# Intrinsic dataset features drive mutational effect prediction by protein language models

**DOI:** 10.64898/2026.03.08.710389

**Authors:** Luiz C. Vieira, Sophia Lin, Claus O. Wilke

## Abstract

Protein language models (pLMs) are commonly used for predicting protein fitness landscapes, but their wide range of performance across datasets remains poorly understood. We evaluated supervised transfer learning on 41 viral and 33 cellular deep-mutational-scanning (DMS) datasets using embeddings from multiple pLMs. We observed consistently lower predictive performance on viral datasets compared to cellular datasets, independent of model architecture or transfer learning strategy. Surprisingly, a simple baseline model that predicted site mean fitness matched or outperformed supervised models on many datasets, highlighting the dominant role of site effects. Analysis of site variability using two metrics, relative variability of site means (RVSM) and fraction of highly variable sites (FHVS), revealed that patterns of fitness variation within and among sites constrain model performance and largely explain the observed differences between viral and cellular datasets. Moreover, splitting training and test data by site, rather than pooling, revealed that supervised models often rely on site effects rather than capturing broader mutational patterns. These findings highlight limitations of current pLMs for mutational effect prediction and suggest that dataset composition, rather than model architecture or training, is the primary driver of predictive success.

**Significance Statement:** Mutational effects prediction with protein language models tends to vary widely in prediction accuracy, depending on the dataset considered. While poor performance is commonly equated with poor model quality, we show here that intrinsic dataset features, such as the variability of fitness values within and among sites, are critical predictors of model performance. Moreover, we show that many existing benchmarks overestimate model performance, by allowing training data to leak into the test set. In fact, in many cases, protein language models barely outperform a naive predictor relying entirely on mean fitness values at individual sites. In aggregate, our study reveals that protein language models are not as reliable for predicting mutational effects as is commonly thought.

---

Protein language models (pLMs) are neural networks trained on large collections of protein sequences using self supervised learning objectives (1–4), inspired by methods originally developed for natural language processing (5). During pretraining, these models learn internal representations, referred to as embeddings, that encode information about protein biochemistry, structure, and evolution (6, 7). These embeddings capture the properties of individual amino acids as well as higher order relationships that reflect local sequence context and global structural and functional organization of proteins (8). Because this information is learned without task-specific labels, a single pretrained model can be reused across different biological problems through transfer learning (9–12). Using this approach, protein language models have been successfully applied to predict protein function (13, 14), structure (15–18), protein interactions (19–21), and the effects of mutations (22–26), establishing them as a core methodology in modern computational biology.

For mutational effect prediction, pLMs are routinely tested on deep mutational scanning (DMS) data, an experimental technique that systematically measures the functional effects of hundreds to thousands of mutations across a protein (27). DMS datasets provide high-resolution maps of how individual amino acid substitutions affect protein stability, activity, and interactions (28–30), capturing the functional consequences of sequence variation in a quantitative, systematic manner that machine-learning models can exploit to learn biologically meaningful representations. However, despite their broad success otherwise, pLMs show widely inconsistent performance on DMS data. While pLMs succeed spectacularly on some DMS datasets they perform poorly on others (23, 31–34). Moreover, there seems to be a systematic trend of lower performance on viral datasets than on cellular datasets (35–37). The causes underlying this substantial variation in model performance are poorly understood.

Here, we systematically evaluate supervised transfer learning across a large collection of viral and cellular DMS datasets. We consider multiple models, transfer learning strategies, and data splitting schemes. By introducing two simple dataset metrics that describe how fitness variation is distributed across and within sites, we show that much of the apparent predictive performance of pLMs is driven by site mutational effect variability, rather than by the predictive capability of the model. Our results demonstrate that viral proteins pose a distinct challenge due to limited within-site variability, that a commonly used splitting strategy inflates performance through data leakage of site information, and that our dataset metrics can predict model success across dataset benchmarks. Together, these findings clarify why pLMs struggle to predict mutational effects in some protein datasets and point toward more rigorous evaluation strategies needed to assess true model generalization.

## Results

### Protein language models do not perform well on viral datasets

The performance of current protein language models (pLMs) in predicting mutational effects from DMS data is highly variable. One systematic pattern is notable, however: Performance appears to be systematically worse for DMS data of viral proteins, as compared to cellular proteins (35–37). To further explore this pattern, we compiled 41 viral and 33 cellular DMS datasets containing only single point mutations and applied supervised transfer learning to predict mutational effects. For each dataset, we separated mutations into training and test sets using pooled splits, whereby all mutations are treated as a single pool and randomly partitioned into either category. This data splitting procedure has been widely used in prior studies (4, 35, 38–44). To predict mutational effects, we performed Lasso regression on mean embeddings extracted from the ESM-2 650 million parameter model (8), evaluating model performance on the held-out test set. We observed that model performance was systematically lower for viral proteins, in particular for longer proteins (Figure 1A, B). A similar trend was observed when replacing ESM-2 with a more recent protein language model, ESM C (45), except performance on viral proteins was even worse (Figure 1C, Supplementary Figure S1).

**Fig. 1.**
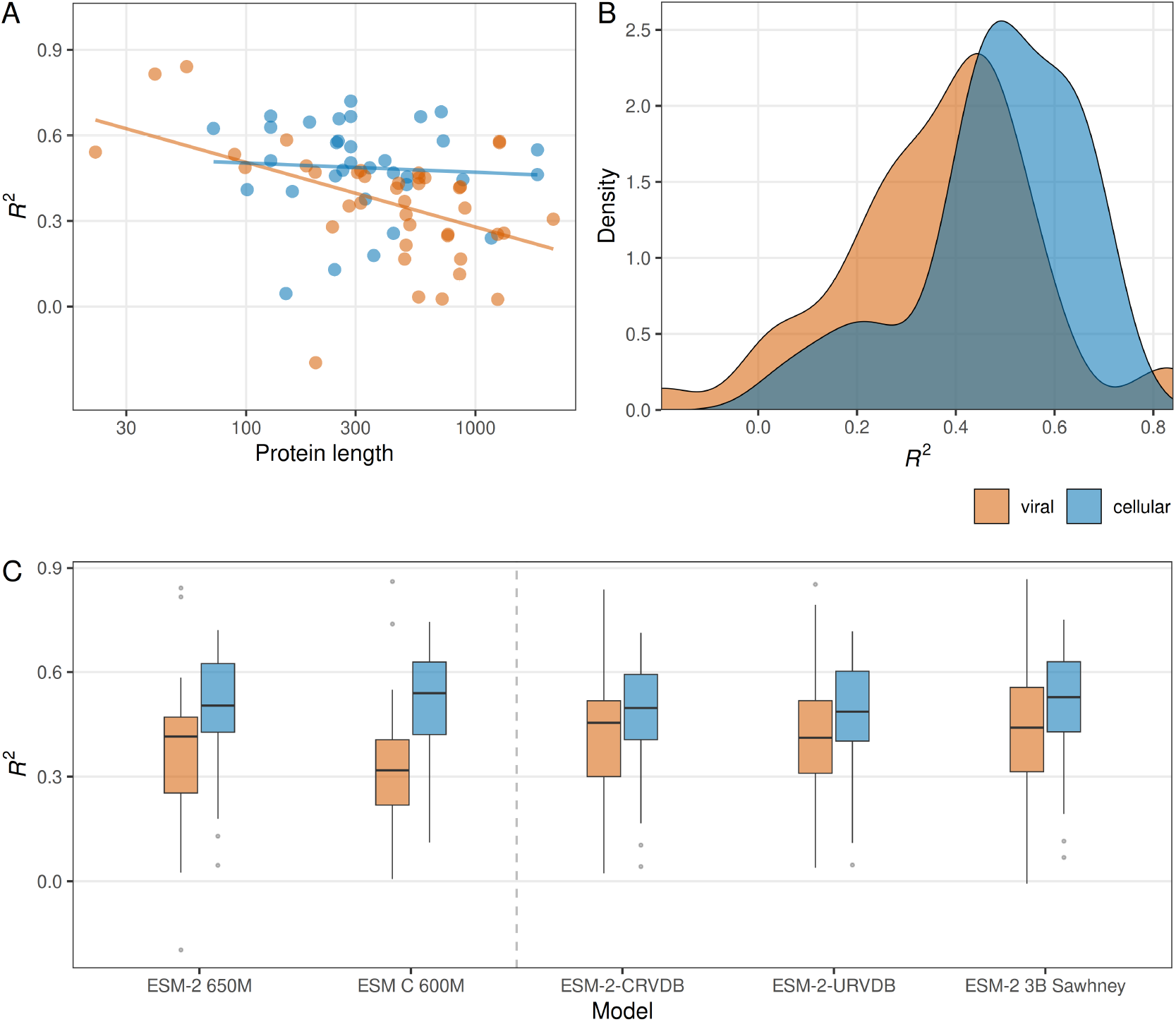
Performance of mutational effect prediction via transfer learning as a function of the dataset type. (A) Mean *R*^2^ under five independent train–test splits are shown as a function of protein length, for both viral and cellular proteins. Each dot represents one DMS dataset. On average, prediction performance tends to be lower for longer proteins. In addition, prediction performance tends to be lower for viral proteins than for cellular proteins. Data were split into training and test sets using pooled splitting. Distributions of the *R*^2^ values shown in Panel A. Prediction performance on viral proteins is consistently lower than on cellular proteins (t-test, *P* = 0.006). (C) Prediction performance for different protein language models. ESM-2 650M and ESM C 600M are the non-domain-adapted base models. The other three models have been finetuned for viral datasets. For both ESM-2 650M and ESM C 600M, prediction performance is significantly reduced for viral proteins compared to cellular proteins (t-test, *P* = 0.006 and *P* = 0.00001, respectively). For the domain-adapted models, there remains a reduction in prediction performance for viral proteins but it is no longer significant (t-test, *P >* 0.05 in all cases). Moreover, ESM-2 CRVDB on viral data showed significantly reduced performance compared to ESM-2 650M on cellular data (t-test, *P* = 0.0493) but ESM-2 CRVDB and ESM-2 3B Sawhney did not (t-test, *P* = 0.05948 and *P* = 0.2157, respectively).

One possible explanation for the reduced performance of pLMs for viral proteins is the underrepresentation of such proteins in the pretraining data. It has been suggested that this limitation could be mitigated by finetuning current models on larger datasets containing more viral sequences and/or using larger models (36). (The process of finetuning on datasets containing specific types of sequences is also referred to as domain adaptation.) To investigate these proposed solutions, we created two finetuned versions of the ESM-2 650 million parameter model. We curated viral protein sequences from the Reference Virus Database (RVDB v30) (46) and created two different subsets of the RVDB dataset, clustered at 80% sequence identity (658,064 sequences) and unclustered (2,696,018 sequences after removing duplicate sequences). We then finetuned ESM-2 650M using masked language modeling, which resulted in our finetuned models ESM-2 650M CRVDB (finetuned on the clustered sequences) and ESM-2 650M URVDB (finetuned on the unclustered sequences). In addition to our own finetuned models, we also investigated a publicly available version of the ESM-2 3 billion parameter model finetuned on viral data (ESM-2 3B Sawhney) (47). The Sawhney ESM-2 3B model has been finetuned using 345,261 sequences obtained from the Virus Orthologous Groups Database (VOGDB) (48).

We found that domain adaptation did not fully close the gap in performance between viral and cellular datasets (Figure 1C). Overall, we observed that when finetuning ESM-2 with viral sequence data the predictive performance for viral proteins somewhat improved, and the predictive performance for celluar proteins somewhat declined, to the point that the difference was no longer statistically significant (Figure 1C). However, for all three finetuned models, performance on cellular datasets remained somewhat higher, on average, than performance on viral datasets. Notably, all three finetuned models showed approximately the same performance. Neither increasing the number of viral protein sequences used for training nor working with the larger ESM-2 3B model made much of a difference.

### Supervised transfer learning is heavily dependent on site variability

Irrespective of the differences in average model performance on viral and cellular datasets, we also observed substantial differences in performance within each group of datasets. Therefore, we reasoned that if we could identify the causes of performance variation within one group of datasets this might also shed light on performance differences between viral and cellular datasets.

We first aimed to establish a minimum baseline of model performance for a given dataset, by considering a simple site-means approach that predicted any unseen mutational effects simply as the average of the known fitness effects at each site in the protein. Surprisingly, for most viral datasets, this site-means method outperformed the pLMs approach (Figure 2A and Supplementary Figure S2A). For cellular datasets, the pLMs did consistently perform better than the baseline method, but only by a small amount (Figure 2A and Supplementary Figure S2A). Moreover, although domain adaptation of pLMs improved performance on viral datasets relative to cellular datasets, it still did not outperform the simple baseline method (Figure 2B and Supplementary Figure S2B-C). This observation highlights that supervised transfer learning with pLMs frequently provides only a minor advantage over naive site averages, and for viral datasets can even be detrimental.

**Fig. 2.**
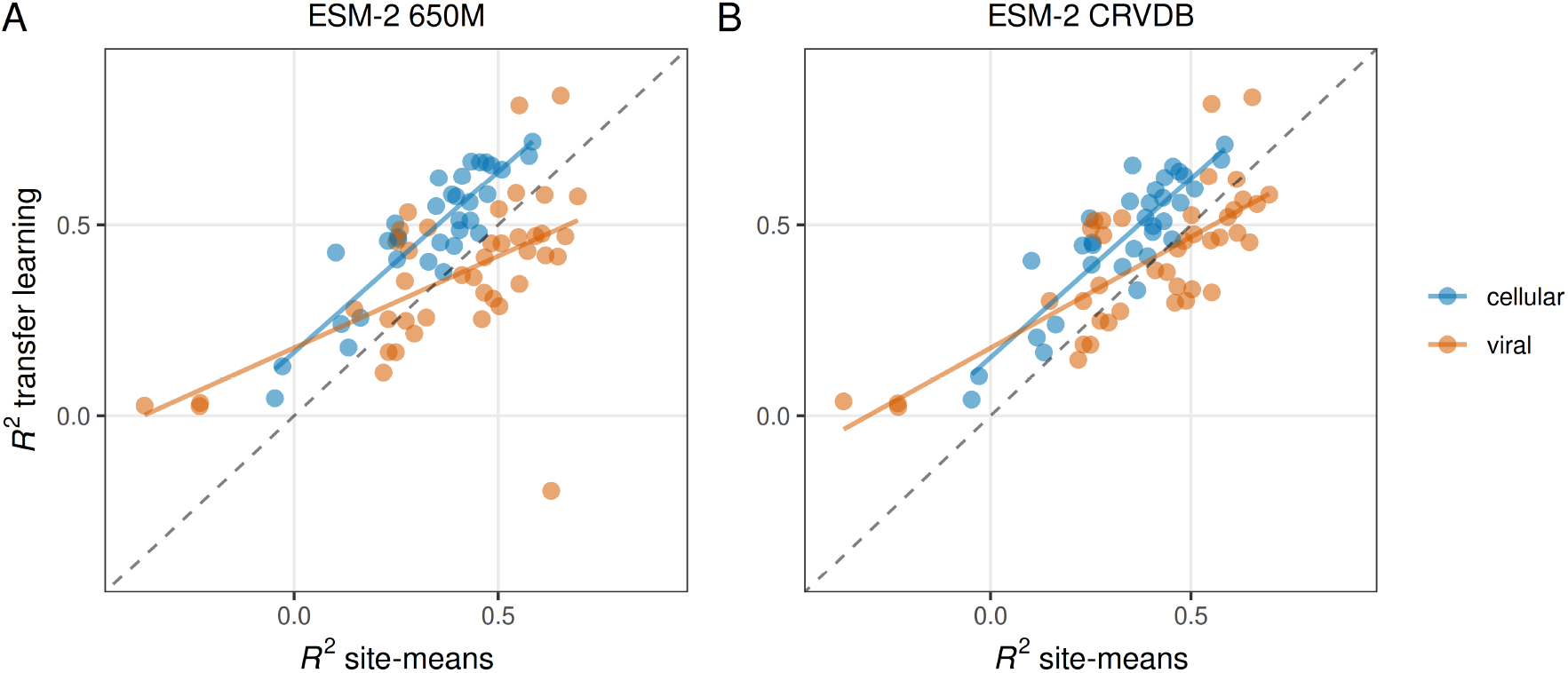
Strong site effects are seen across most datasets. The *R*^2^ achieved under transfer learning (Lasso regression, pooled data splits) is very similar to the *R*^2^ obtained from simply taking the mean at each site in the training data and using it as the prediction for the test data. The dashed line indicates the one–one line where the transfer learning strategy and the site-means strategy perform exactly equally. For cellular proteins, transfer learning performs slightly better than site means (blue dots are all located above the dashed line). For viral proteins, in many cases the site means strategy outperforms transfer learning. (A) Results for the base model ESM-2 650M. (B) Results for the domain-adapted model ESM-2 CRVDB.

Because performance of the site-means approach also varied widely among datasets, we next examined the distribution of fitness effects within and among sites. Given that site identity alone explains over 50% of the variation in many datasets, we hypothesized that the source of poor performance may be intrinsic to the data, particularly to how fitness variability is distributed across mutated sites of a protein.

For this analysis, we defined two complementary measures of site variability. The first, relative variability of site means (RVSM), captures the extent by which site means vary relative to the total variance in the dataset. It is defined as the standard deviation of the site means divided by the overall standard deviation of all data across all sites [Eq. (2)]. Low RVSM values indicate that most of the variability in fitness measurements arises within sites rather than across them, implying that site identity carries limited predictive information. By contrast, high RVSM values indicate that differences between site means account for a substantial portion of the total variation in the data, implying that site effects dominate the overall distribution of fitness effects. The second method, fraction of highly variable sites (FHVS), reflects how many sites exhibit meaningful within-site variation. For every site, we calculate its within-site standard deviation, normalized by the overall standard deviation across all sites. We then determine the fraction of sites for which this normalized standard deviation exceeds 0.7, that is, for which the within-site variation is comparable in magnitude to the overall variation [Eq. (3)].(Note that our results are not strongly dependent on the specific threshold value chosen, see below.) A low FHVS implies that most sites in a protein are not sensitive to mutation, i.e., fitness does not change much when these sites are mutated, whereas a high fraction indicates broader sensitivity to mutation across sites. Together, these two metrics provide a detailed view of how fitness variation is structured across sites.

Our results revealed a clear distinction between viral and cellular datasets in both the RVSM and the FHVS metrics (Supplementary Figure S3). Viral datasets exhibited bimodal distributions for both metrics. For RVSM, a distinct valley around 0.7 separated datasets with low across-site variability (left mode) from those with high variability (right mode) (Supplementary Figure S3A). By contrast, most cellular datasets displayed a more uniform distribution centered around 0.7. The FHVS analysis revealed that a good portion of the viral protein datasets exhibited a low fraction of highly variable sites (FHVS < 0.25). By contrast, none of the cellular datasets had very low FHVS (Supplementary Figure S3B). These results were largely unchanged when using threshold values between 0.3 and 0.8 to define highly variable sites (Supplementary Figure S4).

Next, we quantified the relationship between model performance (measured by *R*^2^) and RVSM. We observed a strong positive correlation between RVSM and prediction performance in cellular datasets and a more moderate correlation in viral datasets (Figure 3A). In general, datasets with higher across-site variability (↑RVSM) achieved stronger fitness predictability, indicating that models with higher *R*^2^ values relied heavily on site differences to make accurate predictions (Figure 3A). However, while RVSM helped explain one of the main sources of predictability in these datasets, it did not account for why performance was poor in some datasets with high RVSM, and in particular viral datasets (Figure 3A).

**Fig. 3.**
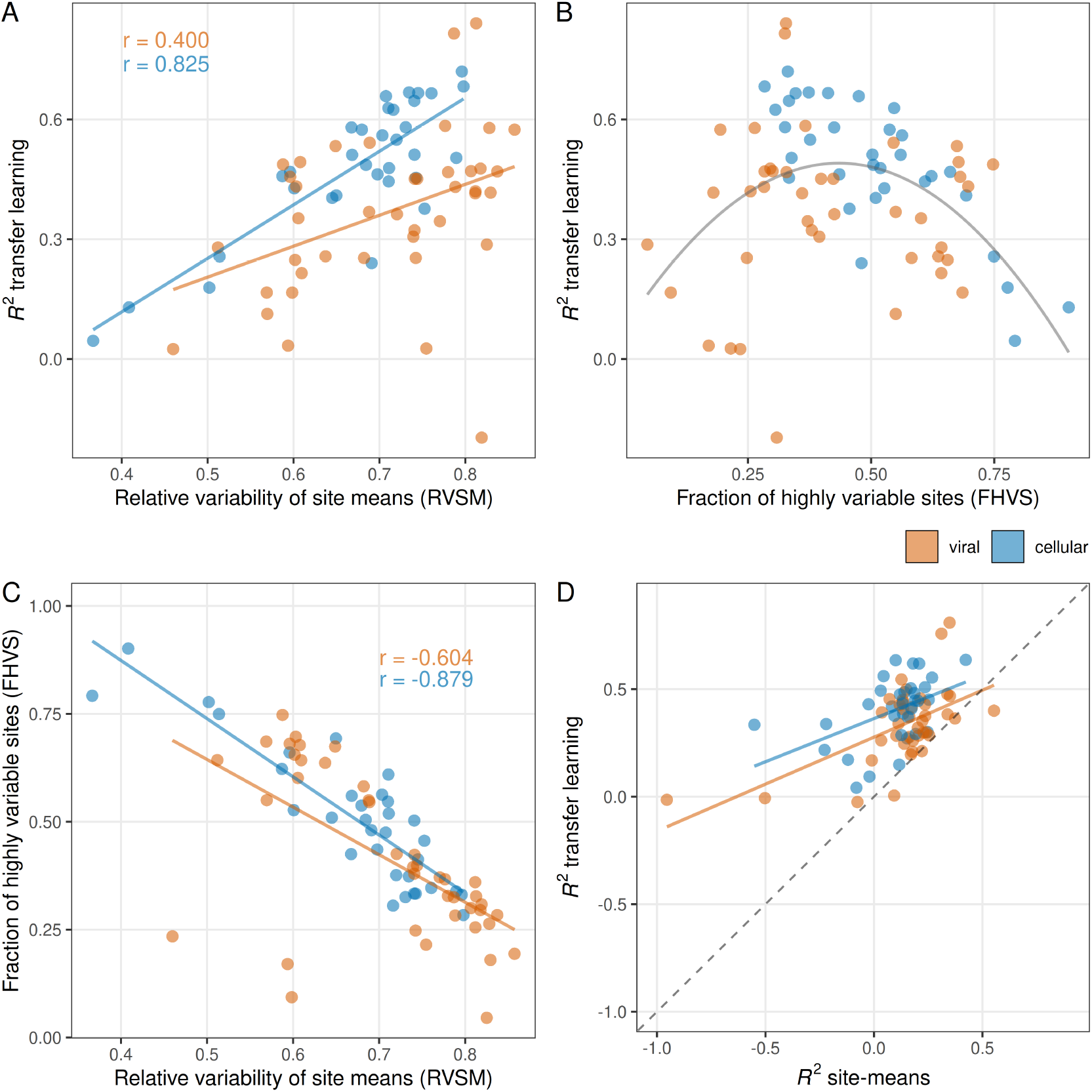
Effect of site fitness variability on transfer learning performance. (A) Transfer learning performance (ESM-2 650M, Lasso regression, pooled data splits) increases with the relative variability of site means (RVSM) in the data. (B) Transfer learning performance is highest at intermediate levels of FHVS. The gray line represents a quadratic fit to the data. (C) The fraction of highly variable sites (FHVS) is negatively correlated to the RVSM. In datasets with more variation among site means, we see less within-site variability. Notably, on average, viral datasets have higher RVSM and lower FHVS. (D) When focusing only on the highly variable sites in each dataset, transfer learning via ESM-2 and Lasso regression consistently outperforms site means for cellular and viral proteins. Moreover, the average performance for these two groups of datasets is nearly identical when focusing only on highly variable sites.

We next investigated the effect of FHVS. Our hypothesis was that higher FHVS would correlate with higher model performance, as models have little to predict when there is no variability within sites. However, instead we found that there was a maximum at intermediate FHVS values (Figure 3B). Models generally performed poorly when FHVS was either very low or very high, and they performed well for FHVS values in the range of 0.25 to 0.5. Also, notably, the lowest FHVS values were only observed for viral datasets, and the highest FHVS values were only observed for cellular datasets. That model performance is maximized at intermediate FHVS values can be explained by the observation that RVSM and FHVS are strongly negatively correlated (Figure 3C). Thus, while model performance generally tends to increase as either quantity increases, the two quantities trade off with each other and no dataset has both high RVSM and high FHVS. These results suggest that achieving reasonable predictive performance under pooled data splits requires a balance between variability within sites and across sites.

Because viral and cellular datasets were so clearly differentiated by the fraction of highly variable sites, with many viral datasets displaying a distinct lack of such sites (Figure 3B), we hypothesized that the low performance of pLMs on these datasets could be caused by an excess of these uninformative sites, where mutations have fundamentally no effect on the measured fitness of the protein. To test this hypothesis, we retained only the highly variable sites in all datasets and discarded all other sites before refitting the models. After this adjustment, transfer learning with ESM-2 consistently outperformed the site-means model (Figure 3D). Moreover, the performance gap between viral and cellular proteins was virtually eliminated. As before, this result was not strongly dependent on the specific threshold choice in the FHVS calculation (Supplementary Figure S5). Thus, we conclude that the main difference between viral and cellular datasets is the relative lack of highly variable sites in viral data, and this lack is the primary cause of the differences in model performance between viral and cellular datasets.

### Supervised transfer learning fails to generalize under site-stratified splits

Having shown that supervised transfer learning models rely strongly on site variation, we next examined how data splitting strategies affected regression performance on DMS datasets. Prior works have typically used pooled splits, which allocate mutations from the same sites across both training and test sets. As we established above, this approach inflates performance due to data leakage, because the model has access to the same site information during training and test. To address this issue, we compared the commonly used pooled split with a more robust site-stratified split that prevents data leakage (Figure 4A). In this second strategy, all mutations at a given site are assigned exclusively to either the training or the test set. This design prevents the model from learning site-specific average effects and instead requires it to generalize to entirely unseen sites.

**Fig. 4.**
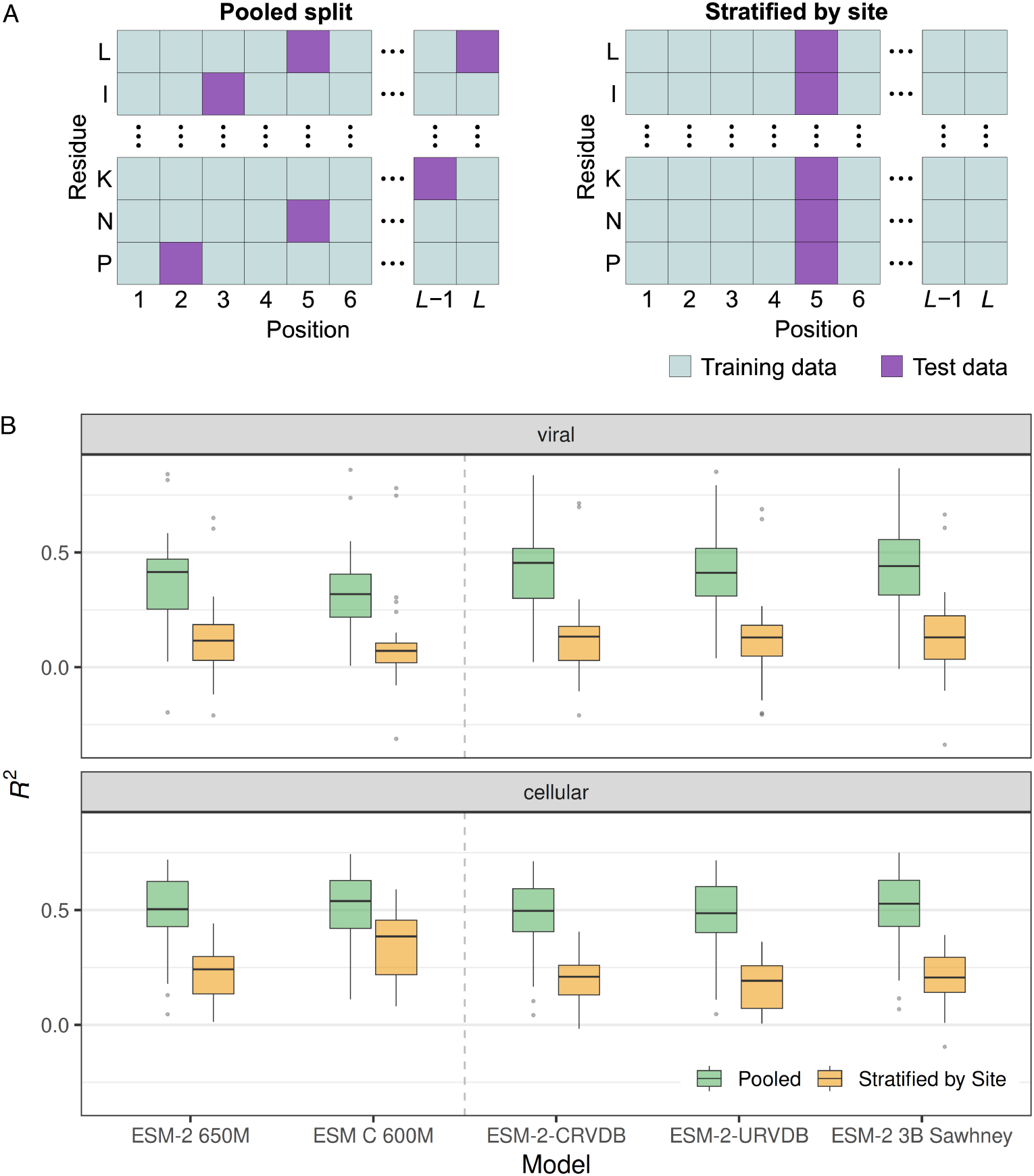
Transfer learning performance is substantially affected by the data splitting strategy. (A) Schematic representation of data splitting strategies. Pooled split: Mutations from the same site can appear in both training and test sets, allowing the model to see site-specific information during training. Site-stratified split: All mutations from a given site are assigned exclusively to either the training or the test set, ensuring the model must generalize to unseen sites. (B) Performance of transfer learning by different splitting strategies. For both viral and cellular datasets, and for all protein language models considered, the *R*^2^ obtained from transfer learning (Lasso regression) is significantly higher when data is split according to the pooled strategy as compared to splits stratified by site. (Significance was assessed by t-tests comparing pooled *R*^2^ vs. stratified-by-site *R*^2^ separately for each model and dataset type, *P* < 0.005 for all comparisons.)

We found that splitting the data by site lead to a substantial decrease in performance for all tested models, with no substantive difference between viral and cellular datasets, both of which were similarly affected (Figure 4B). On cellular datasets, however, ESM C consistently performed best even under site-stratified splits, followed by its predecessor model ESM-2. This result indicates that architectural improvements in a model can enhance generalization.

We also asked whether the relationships between model performance and either RVSM or FHVS remained visible after switching from pooled splits to splits stratified by site. We found that indeed this was the case (Supplementary Figure S6). While the strength of the relationship was attenuated in both cases, *R*^2^ continued to increase with increasing RVSM even under splits stratified by site (Supplementary Figure S6A), and it tended to reach a maximum at intermediate values of FHVS (Supplementary Figure S6B). This finding highlights that the usefulness of RVSM and FHVS as predictors of model performance is not simply due to data leakage from training to test under pooled splits. Instead, RVSM and FHVS are genuine predictors of model performance, all else being equal. We expect that most models will struggle with datasets that have low RVSM and/or either low or high FHVS.

### Transfer-learning results are qualitatively unchanged under finetuning

All transfer-learning results reported so far were obtained under a two-step process of first extracting embeddings from a pLM and then using them as features in Lasso regression. It is known that this process typically results in lower model performance than a strategy that includes finetuning the pLM (35, 49). Therefore, it is reasonable to ask whether results would look substantially different if we had used fintetuning instead.

To evaluate the finetuning approach, we added a regression head to ESM-2 650M and then finetuned the model via low-rank adaptation (LoRA). As expected, we found that the *R*^2^ values of the finetuned models were consistently higher than those obtained under Lasso regression, regardless of the data splitting strategy employed (Supplementary Figure S7). However, we also saw that finetuning did not qualitatively change our overall observations. First, most importantly, *R*^2^ values obtained with Lasso regression and *R*^2^ values obtained with finetuning were strongly correlated, and the improvement gained from finetuning was generally small. Finetuning did not help much for datasets where Lasso regression performed poorly. Finetuning also did not help much for training/test splits stratified by site. Finally, the relative poorer performance on viral proteins as compared to cellular proteins remained during finetuning.

In summary, while finetuning is a viable strategy to obtain maximum predictive performance from a protein language model, it cannot overcome inherent limitations of insufficient variation within and among sites in the dataset, nor can it overcome limitations of the model not generalizing well when training/test splits are stratified by site.

### Variability metrics reliably predict model success across ProteinGym datasets

Because our analysis up to this point was limited to ESM-style models, we wanted to investigate whether similar results could be observed for a wider class of models. To this end, we turned to the ProteinGym project (50), which systematically benchmarks a wide range of different models against an array of DMS datasets. Specifically, we asked two questions: First, is the strong impact of the data splitting strategy on model performance visible in the ProteinGym results? Second, can model performance reported by ProteinGym be predicted by the two variability measures we have introduced, RVSM and FHVS? If other models also mostly memorize mean mutational effects at sites and reproduce them for the held-out mutations in the training set, then pooled splits should show elevated model performance compared to more rigorous splitting strategies. More generally, model performance should be influenced by RVSM and FHVS, regardless of the model considered.

We restricted our analysis to single-mutant DMS datasets available in ProteinGym to focus specifically on site effects. ProteinGym reports the Spearman correlation coefficient *ρ* for each DMS dataset under three data-splitting schemes used for five-fold cross validation: (i) pooled splits, referred to as “random” in ProteinGym, in which individual mutations are randomly assigned to one of the five folds; (ii) “modulo”, a version of stratified by site, in which mutated sites are assigned to folds using the modulo operator (site 1 is assigned to fold 1, site 2 to fold 2, and so on, returning to fold 1 at position 6 and continuing this pattern along the sequence); and (iii) “contiguous”, also a version of stratified by site, in which the sequence is divided into five consecutive segments and mutations are assigned to folds based on the segment in which their position falls (51).

Among the models benchmarked by ProteinGym, we focused on five top-performing supervised models that rely on pLM embeddings as protein representation. ProteinGym evaluates both non-augmented and augmented supervised models. The top-scoring non-augmented models include Kermut (52), a Gaussian process regression model with a composite kernel that captures mutation similarity and provides uncertainty estimates, using mean-pooled embeddings from the 650M-parameter ESM-2 model as input, and ProteinNPT (51), a semi-supervised pseudo-generative model that jointly represents protein sequences and property labels, learning their interactions for supervised prediction. The top-scoring augmented models are MSA Transformer (53), Tranception (54), and ESM-1v (23). For each of these models, a separate ridge regression is trained on mean-pooled embeddings combined with a sequence density score, defined as the inferred log-likelihood under a pretrained evolutionary probability model. This strategy combines model-specific pretrained embeddings with evolutionary information to improve supervised prediction (55). We found that all models performed best under the pooled split strategy, followed by the modulo strategy and then the contiguous splits, respectively (Figure 5A). Additionally, the non-augmented models outperformed the augmented models (Figure 5A). Together, these results corroborate our hypothesis that whenever the pooled split is used as the primary data-splitting strategy, it contributes to information leakage during modeling and inflates prediction accuracy.

**Fig. 5.**
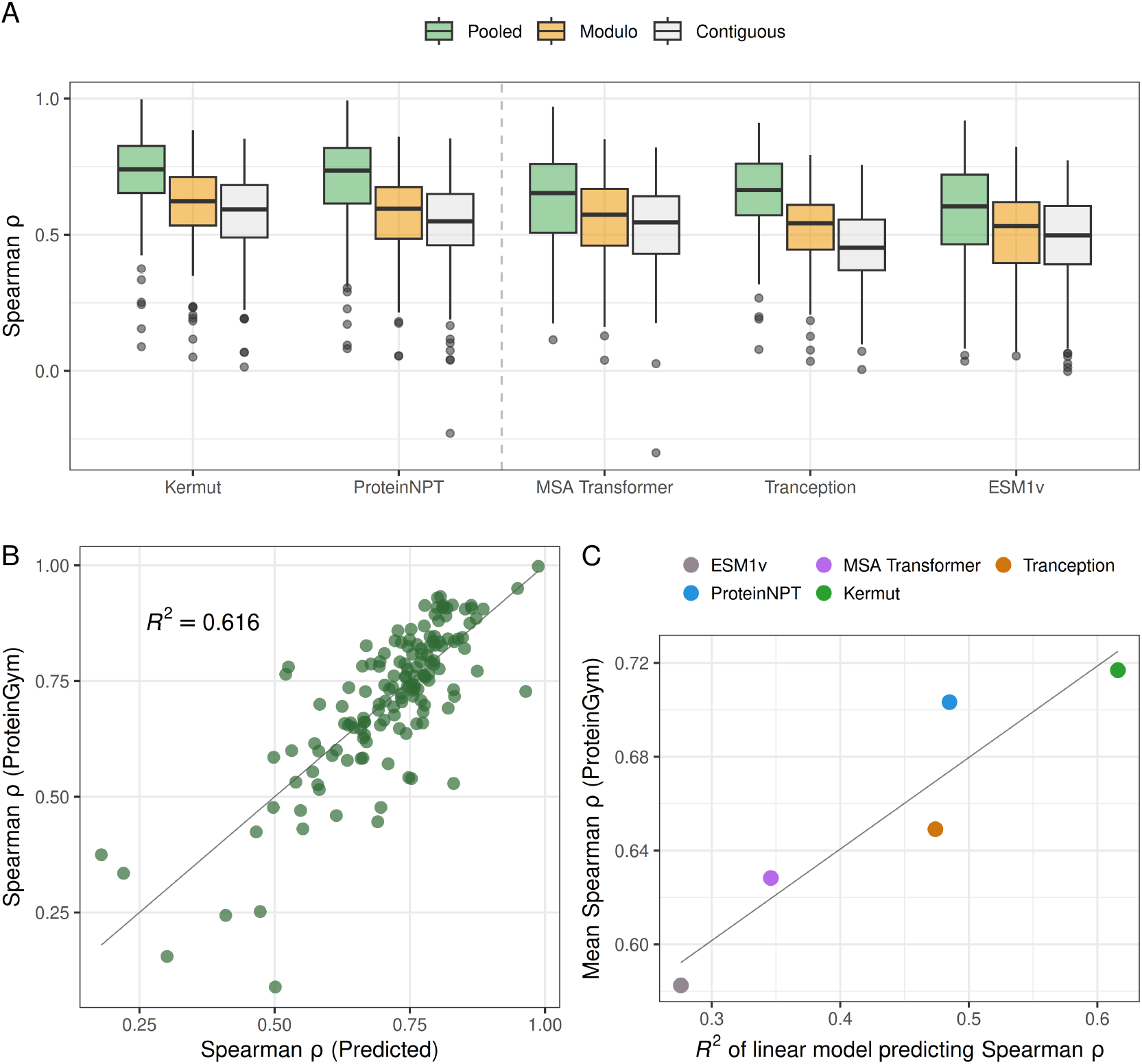
Dataset splitting strategy and fitness variability within and among sites impact model performance reported by ProteinGym. (A) Model performance in ProteinGym (as measured by Spearman *ρ*) is consistently better for pooled splits than for splits stratified by site. Note that ProteinGym has two stratified splitting strategies, “modulo” where test sites are distributed uniformly throughout the protein and “contiguous” where a contiguous region of the protein is held out for testing. Models consistently perform the worst on modulo and contiguous splits. For every model, the pairwise comparisons between pooled and either modulo or contiguous splits were significant according to Tukey’s test, with *P* < 0.01 in all cases (Supplementary table 1). The dashed line separates non-augmented models to the left from augmented models to the right. (B) Model performance reported by ProteinGym for pooled splits can be predicted by a simple regression model using site-variability metrics RVSM and FHVS as predictors. Each dot represents one ProteinGym dataset, and results are shown for the Kermut model, the overall best performing model in ProteinGym. (C) Average model performance in ProteinGym correlates with how well site variability measures predict model performance. The higher the *R*^2^ in the linear model, as shown in Panel B for Kermut, the higher the mean Spearman *ρ* reported by ProteinGym.

To examine the relationship between model performance and RVSM and FHVS, we regressed the Spearman correlation coefficient *ρ* reported for each model and dataset against the predictor variables RVSM and FHVS calculated for each dataset. In all cases, we used the *ρ* values reported for pooled splits. For the top-performing supervised model, Kermut (52), the two variability metrics explained a substantial fraction of the observed performance variation, with an *R*^2^ of 61% (Figure 5B). A similar pattern was observed for the next four best-performing models, with explained variance ranging from 27% to 40% (Supplementary Figure S8). We further found a strong linear relationship between the *R*^2^ in our regression model predicting ProteinGym *ρ* and the mean ProteinGym *ρ*, indicating that the better a model performs the more its performance is predicted by site variability patterns (Figure 5C). Together, these results suggest that performance differences across datasets are largely driven by positional fitness variability, indicating that supervised models may rely more on site effects than on learning complex mutational interactions.

**Table 1.**
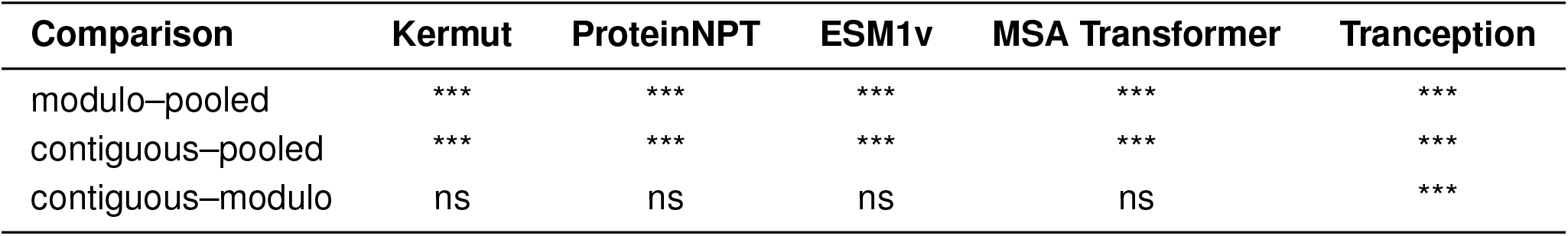
Statistical comparison of model performance across different train–test split strategies on the ProteinGym benchmark. For each model, a one-way ANOVA was performed using Spearman correlation across splits, followed by a Tukey post hoc test for pairwise comparisons between split strategies. The “Comparison” column indicates the pair of split strategies being compared (modulo, contiguous, and pooled). Each subsequent column reports the significance level of the performance difference for the corresponding model (Kermut, ProteinNPT, ESM1v, MSA Transformer, and Tranception). Significance is denoted as: *** *P* < 0.001, ** *P* < 0.01, * *P* < 0.05, and ns, not significant.

### Supervised transfer learning fails to generalize from single to double mutants

All our work so far has focused on single point mutations, but problems of inappropriate train/test splits and data leakage can also occur when trying to predict multi-mutant fitness. To explore this application area, we collected 58 double-mutant DMS datasets and evaluated three distinct modeling approaches. First, we performed a pooled split, where all variants were randomly divided into training and test sets, and trained the regression models on ESM-2 650M and ESM C 600M embeddings. Second, we trained exclusively on single mutants and evaluated the models on double mutants using the same embedding-based regression framework. Third, we implemented an additive baseline model trained on single mutants and evaluated on double mutants, where the predicted fitness of a double mutant was estimated as the sum of the corresponding single-mutant fitness effects.

We found that the pooled split produced high *R*^2^ values for both regression models (Supplementary Figure S9A). By contrast, when models trained exclusively on single mutants were used to predict double mutants, performance was poor, even though ESM C 600M performed somewhat better than ESM-2 650M (Supplementary Figure S9A). These results suggest that high performance under the pooled split is at least in part due to data leakage. The protein language models cannot on their own generalize from single to double mutants and correctly predict the fitness of double mutants purely from knowing the fitness of single mutants.

However, in comparison to the simple additive baseline, the ESM C embedding-based model did indeed perform better on the vast majority of datasets (Supplementary Figure S9B), even if in absolute terms performance was poor. This result indicates that protein language models do have some knowledge about interactions between sites, but for current models this knowledge is not sufficient to generalize fitness predictions from single to double mutations.

## Discussion

We have found that current protein language models (pLMs) show highly variable performance in mutational effect prediction across datasets, and in particular between viral and cellular datasets. Moreover, neither domain-adapting the base pLM nor finetuning a pLM on task-specific mutation data consistently improves predictive accuracy, suggesting fundamental limits imposed by the structure of the underlying fitness data. By comparing pLM performance to a naive predictor using only site means, we have found that site effects frequently dominate, with the naive site-means predictor matching or outperforming supervised transfer learning models in many cases. By introducing two informative summary statistics, the relative variability of site means (RVSM) and the fraction of highly variable sites (FHVS), we have shown that high among-site variation is a key determinant of dataset predictability, and that apparent model success is often driven by differences across sites rather than by learning substitution-specific effects. We have further demonstrated that commonly used pooled data splits tend to inflate performance estimates due to data leakage. Together, these results indicate that much of the predictive signal attributed to supervised transfer learning arises from simple positional effects.

Our results suggest that the entire field of mutational effects prediction with pLMs may be over-estimating the performance of their models. Under pooled data splits, which are widely used (4, 35, 38, 56–64), datasets with low within-site variability become artificially predictable. This has significant implications for applying these models to protein-engineering tasks, where practitioners often need to predict effects at positions with limited experimental data (65). When site identity explains most of the variance, even sophisticated embedding-based models primarily learn to recapitulate site means rather than capturing genuine sequence–function relationships. Good model performance in this case does not indicate that the embeddings encode deeper biochemical or evolutionary information. Instead, it reflects data leakage of site effects into the test set, which inflates model performance. By stratifying training/test splits by site, this shortcut is removed, forcing models to rely on features that generalize across sequence context rather than memorizing site identity.

Moreover, even though we have focused on pLMs here, we believe our main findings, that pooled splits are responsible for inflated performance figures, apply more broadly. In fact, we found similar results for all the top performing models in the ProteinGym benchmark, regardless of model type. Consistent with our interpretation, other studies that explicitly evaluated various models under different stratification approaches reported substantial performance drops compared to pooled splits (66–69), highlighting the extent to which position identity drives apparent accuracy. In this context, we would like to highlight that the most widely used benchmarking study for fitness effects prediction, the ProteinGym, has potentially contributed to confusion on this topic, by writing (51): “We note that there is no inherent issue with using a Random [i.e., pooled] cross-validation scheme to estimate the performance of predictive models. However, the conclusions drawn and the generalizability claims based on it require careful consideration.”

The ProteinGym data provides one additional insight. When comparing their modulo strategy (which spreads out test sites uniformly across the protein sequence) to their contiguous strategy (which takes contiguous stretches of positions as either training or test data), we have seen additional performance degradation, if minor, under the latter. This observation suggests that there may be autocorrelation among fitness values between sites, such that data leakage can be a problem not only when mutations belonging to the same site are present in both the training and the test set but also when among neighboring sites some are in the training set and others in the test set.

We have also investigated to what extent current embedding-based approaches can generalize from single-mutant measurements to double variants. Although pLMs are expected to encode contextual and structural information (70), our results suggest these representations alone are insufficient to accurately capture higher-order epistatic interactions. In most datasets, models trained on single mutants failed to provide good predictions for double mutants, even if they outperformed a simple additive baseline model. This observation highlights a broader challenge: while pLM embeddings capture local mutational tolerance and sequence context, epistasis emerges from complex residue interactions involving long-range structural contacts, conformational dynamics, or functional constraints (71–73) not fully represented in current embeddings. Improving combinatorial variant prediction will likely require training on higher-order mutation data, explicit structural or biophysical integration, or model architectures explicitly designed to represent non-additive fitness landscapes. It is also possible that concepts from covariation analysis (74) can be merged with protein language models for improved prediction of epistatic effects. This is beyond the scope of the present work but may be a valuable avenue for future research.

It is interesting to see that viral and cellular datasets for which prediction accuracy is poor seem to fail in different ways. When plotting model performance against FHVS, the viral datasets fall on the left side of the parabola, characterized by low variability within sites, whereas the cellular datasets fall on the right side, characterized by high variability within individual sites but relatively low variability across sites. In both scenarios, these constrained variability patterns reduce the effective signal available for statistical learning.

The divergent pattern observed between viral and cellular proteins may in part be due to inherent differences in the structures of these proteins. In particular, many viral proteins are primarily structural, such as capsid proteins or spike proteins, and their structures are unique to viruses and have no homology in cellular organisms. We have not explored this possibility here. If the unique structures of viral proteins are the cause of reduced performance in predicting DMS data, then it is possible that models explicitly taking advantage of structural data could perform better on these proteins. However, as a counterpoint, we would like to emphasize that pLM embeddings already encode substantial structural information (8, 70), and incorporating explicit structural features appears to yield only modest or no improvements in DMS prediction (75).

Alternatively, the differences between viral and cellular proteins may also reflect differences in their underlying evolutionary paths. Although mutations in DMS experiments are introduced artificially, the measured fitness effects still capture how tolerant each protein is to amino acid changes (27). Viral proteins, shaped by high mutation rates, larger population sizes, short generation times, and strong selection pressures (76), may have evolved a broader mutational tolerance across many sites (77, 78). By contrast, cellular proteins often operate under tighter structural and functional constraints (79), which can concentrate tolerated variation in specific regions while keeping much of the sequence highly conserved. Future work should consider that predictive performance is highest at intermediate within-site and across-site variability. Designing DMS experiments that capture a balanced spectrum of mutational effects may therefore yield more informative data for machine learning models.

An important limitation of DMS datasets is experimental noise. Multiple independent DMS measurements of the same phenotype in the same protein are often only moderately correlated (31, 80), and this experimental noise places an upper bound on how good any machine-learning prediction can be. However, we do not expect this issue to alter the central conclusions of this study. Experimental noise would not be expected to systematically favor one train/test splitting strategy over another, and there is also no good reason to assume that experiments on viral proteins are inherently more noisy than on cellular proteins or vice versa.

Our work here has shown, consistent with previously published work (35, 49), that finetuning pLMs usually leads to superior performance compared to other transfer-learning techniques. However, we observed here that even when pLMs were finetuned, they did not overcome the reduced performance observed on splits stratified by site. This result suggests that finetuning alone is insufficient to address the generalization challenge posed by site stratification. While finetuning enables models to learn more complex non-linear relationships from the data (81), the consistent performance gap on site splits indicates that both finetuned and frozen pLM approaches may share a common limitation in how they process sequence information for prediction. Further investigation into alternative architectures or input representations that better preserve position-specific information may be necessary to improve generalization to unseen mutated sites.

Most pLMs are pretrained on broad sequence databases spanning diverse taxa and protein families (1, 4, 8, 45), whereas downstream applications such as variant effect prediction, fitness estimation, or phenotype prediction typically focus on a restricted subset of proteins. Consequently, adapting the model to sequences from the relevant domain allows the internal representations to better align with the functional signals that matter for the task (82), reducing the gap between the pretraining distribution and the target data (83). This typically leads to more informative embeddings and improved generalization, especially when labeled data are limited. Consistent with this rationale, we observed improved performance after domain adaptation on viral proteins, although the improvements did not fully close the gap to cellular data. In particular, the domain adapted models could still not consistently outperform the site-means model on viral data. Nevertheless, our results highlight that current protein language models, and especially ESM C, appear to have been insufficiently trained on viral sequences, and further improvements in developing models for viral sequence data are possible. At the same time, there are inherent differences in the distribution of fitness effects between viral and cellular proteins that make mutational effects prediction in viral proteins more difficult than in cellular proteins, and it is unlikely that this difference can be overcome entirely by domain-adapting pLMs to viral sequence data.

The poor performance of ESM C on viral datasets is notable, considering its strong performance on cellular proteins compared to other models (35, 45). The most likely explanation for this underperformance is its training data: Although no paper describing ESM C has been published, we note that for the publicly available version of ESM 3 (17), the team omitted viral sequences during model training, citing safety concerns as the reason. It is likely that similar choices were made during ESM C training. Going forward, we would recommend against using ESM C when working on any viral proteins. At the same time, we acknowledge that for cellular proteins ESM C is usually the best performing model by a wide margin.

## Materials and Methods

### Data collection

To evaluate mutational effects prediction, we used existing datasets previously compiled in the literature. Specifically, we used 41 deep mutational scanning (DMS) datasets of viral proteins (34) and 33 datasets of cellular proteins (84). All DMS datasets consisted of single amino acid substitutions with associated measurements, such as fitness, stability changes, or antibody escape, for each mutation. For every dataset, we created a FASTA file containing the mutant protein sequences and a CSV file including both mutant sequences and corresponding target measurements.

For domain adaptation of the ESM-2 650M model to viral sequence data, we downloaded protein sequences from the Reference Virus Database (RVDB v30) (46). We used two versions of the dataset: one clustered at 80% identity (C-RVDB) and one unclustered (U-RVDB). Before training, we filtered the original dataset to remove sequences longer than 1,600 residues and to discard sequences containing non-canonical amino acids (X, B, Z, or J). After filtering, we retained 658,064 and 2,696,018 unique sequences for the clustered and unclustered dataset, respectively. We then randomly subdivided these sequences into 80:10:10 splits for training, validation, and test sets.

The single-to-double experiments were conducted using the ProteinGym (50) dataset, downloaded from the DMS Assays (substitutions) collection available on their website. We preprocessed the data by retaining only single and double mutants, resulting in a total of 58 DMS datasets.

### Domain adaptation

Domain adaptation was performed via masked language modeling, where 15 percent of positions in each sequence were masked. Each masked position had an 80% chance of being replaced with a `<mask>` token, a 10% chance of being replaced with a random token, and a 10% chance of remaining unchanged. We refer to the resulting finetuned models as ESM-2 650M CRVDB and ESM-2 650M URVDB, depending on whether they were trained on the clustered or unclustered versions of the RVDB, respectively.

Training was run on the training set for ten epochs using eight A100 GPUs with a per-GPU batch size of 32. At the end of each epoch, model performance was evaluated with the validation set, and at the end of finetuning the model was evaluated one more time with the test set. We used the AdamW optimizer with a learning rate of 4 ×10−4, weight decay of 1 ×10^−2^, and AdamW hyper-parameters *β*_1_=0.9 and *β*_2_=0.95. The training objective was cross-entropy loss applied to the masked positions. The learning rate schedule included a linear warmup during the first epoch, increasing from 0.1× the base learning rate to the base learning rate by the end of the epoch. For the remaining epochs, the learning rate followed a cosine annealing decay, gradually decreasing to 0.1 × the base learning rate. Distributed data parallelism (DDP) was used for training, with early stopping applied using a patience of five epochs, and the checkpoint with the lowest validation loss was used as the final model.

In our first version of the domain-adapted models, we observed a substantial drop in performance on the cellular proteins. We interpreted this result as catastrophic forgetting (85), a known effect where a model loses performance on a previously learned tasks after being further trained on a new domain. To mitigate this effect, we applied two strategies. First, we added cellular proteins to the training set by selecting all unique reviewed UniProt sequences matching the query `(NOT (taxonomy_id:10239) AND (length:[* TO 1600]) AND (reviewed:true))`, where taxonomy ID 10239 excludes viral sequences. We then clustered this set at 50% sequence identity, resulting in 121,273 sequences added to the training set. Second, we finetuned only the last three layers of the ESM-2 650M model, to limit the risk of overfitting.

### Protein language model embeddings

For transfer learning, we extracted embeddings from protein language models. pLM embeddings are matrices of *n× d* dimensions, where *n* represents the protein sequence length and *d* represents the embedding dimension, which differs for different model variants and generally increases for models with more parameters. Here, we exclusively used embeddings from the respective model’s final layer. Moreover, we used mean pooling (35) throughout, where we averaged embeddings along the sequence-length dimension, resulting in a numeric vector of length *d* as feature vector for downstream modeling.

For ESM-2 model variants, we obtained embeddings using the extract.py script available at the ESM-2 GitHub repository (https://github.com/facebookresearch/esm/blob/main/scripts/extract.py). To extract ESM-2 embeddings of sequences longer than 1022 residues, we modified the script by increasing the max truncation length of sequences to 5000. See the script available at: https://github.com/ziul-bio/ViCAM/blob/main/scripts/extract_esm2.py

For ESM C 600M, we followed the instructions provided on the Evolutionary Scale GitHub page: https://github.com/evolutionaryscale/esm. Based on these instructions, we wrote a custom extraction script, available at: https://github.com/ziul-bio/ViCAM/blob/main/scripts/extract_ESMC.py. The main adjustment we did compared to the published instructions was that after loading the model, we changed the data type to float32 to ensure the reproducibility of results as explain indetails at (35).

For the viral-specific model finetuned by Sawhney et al.(47), the base pretrained ESM-2 model `facebook/esm2_t36_3B_UR50D` and its corresponding tokenizer were loaded using the Hugging Face Transformers library. A parameter-efficient finetuning (PEFT) adapter trained on viral proteins was then loaded from the check-point directory `rsawhney_esm2_3B,` available at https://huggingface.co/rsawhney/finetuning-plms/tree/main/esm2_3B/mlm. Mean pooled embeddings were then extracted using a custom extraction script available at: https://github.com/ziul-bio/ViCAM/blob/main/scripts/extract_esm2_3B_tuned.py.

### Train/test splitting strategies

To split data into training and test sets, we applied two distinct splitting strategies: (1) Under the *pooled* splitting strategy, the train/test categorization happens at the level of individual mutations. This splitting strategy ignores site information and will in general result in the same sites occurring in both the training and the test set. (2) Under the *stratified-by-site* splitting strategy, data splitting happens at the level of individual sites, not individual mutations. In other words, all mutations at the same site are jointly assigned to either the training set or the test set.

For the pooled splitting strategy, we randomly subdivided all mutations in a dataset into 80% for training and 20% for testing, using scikit-learn’s `train_test_split` function. This procedure was repeated *k* times with different random seeds. We used *k* = 5 for Lasso regression and *k* = 3 for model finetuning.

For the *stratified-by-site* strategy, we randomly picked a site to be allocated to the test set, added all mutations at that site to the test set, and repeated with another randomly chosen site until the test set reached 20% of the entire dataset, with the remaining sequences used for training. For example, if a protein had mutations at 10 sites, sites 3 and 9 might be assigned to the test set. If the sum of their sequences exceeds the 20% threshold, sites are resampled until we select sites that correctly populate the desired set size, while the remaining sites are used for training. As before, this procedure was repeated *k* times with different random seeds, with *k* = 5 for Lasso regression and *k* = 3 for model finetuning.

To generate a reproducible random seed for each dataset and replicate, we formed a string by concatenating the overall standard deviation. The site means are computed as dataset name with the replicate index (for example, *BLAT_ECOLX_Ranganathan2015_1*). This string was hashed using the MD5 algorithm, and the first eight hexadecimal characters of the hash were converted to an integer. This integer served as the random seed for that specific dataset–replicate pair. This approach ensured that all methods, whether Lasso regression or finetuning, received identical training and test splits for the same replicate, allowing for fair and direct comparisons across models.

### Regression modeling with Lasso

To predict fitness effects from protein embeddings, we employed LassoCV, a cross validation based version of the Lasso algorithm (86). LassoCV retains the feature selection properties of Lasso by reducing the coefficients of less relevant features to zero while selecting the optimal regularization parameter through cross-validation, thereby identifying the most important predictors. All modeling was conducted in Python using methods from scikit-learn. Before any modeling, the dataset targets were scaled using `StandardScaler`.

To determine the optimal regularization parameter *α* within each round of cross-validation, the LassoCV algorithm as implemented by scikit-learn generates a sequence of 100 *α* values, spanning three orders of magnitude in range. For each *α*, Lasso regression is performed on the training data via 5-fold cross-validation. The optimal *α* is selected based on the lowest cross-validation error, and a final Lasso model is fit using the best *α* and the entire training dataset and evaluated on the test set.

We repeated the above process five times in a second-level validation process, using training/test splits as described in the previous subsection. The final reported model performance was the mean *R*^2^ calculated over all test sets from the five-fold replication.

### Additive modeling for DMS datasets with double mutants

We implemented an additive baseline model using only single-mutant measurements for training and double mutants for evaluation. Predicted fitness values for double mutants were obtained by summing the fitness effects of their corresponding single mutations. When a test-set mutation was not observed among the training single mutants, its effect was approximated using the mean fitness effect of mutations at the same position. If no measurements were available for that position, the median fitness value of the training dataset was used as a fallback estimate.

### Finetuning ESM-2 650M with LoRA

To finetune ESM-2 650M for variant effect prediction, we leveraged PFET (Parameter-Efficient finetuning) (87) to apply LoRA (Low-Rank Adaptation) (88) layers (specifically, `q_proj` and `v_proj`) to the models while freezing all other layers. The LoRA layers were set with a rank of 8, a scaling factor of 64 (`lora_alpha`), and a dropout rate of 0.1. We replaced the language-modeling head with a regression head implemented in a pre-norm style. The head takes as input the model’s embedding vector (dimension 1280) at the BOS token. The embedding is first normalized with `LayerNorm`, then passed through a linear projection of the same dimensionality as the embedding size, followed by a GELU activation and dropout (rate 0.1). A final linear projection maps this hidden representation to a single regression output.

We selected 42 DMS datasets (15 cellular and 27 viral) for finetuning, restricting to single point mutation datasets with more than 2,000 observatioms and proteins shorter than 1,300 residues. Each dataset was split into training and test sets using an 80:20 ratio. Models were trained for up to 50 epochs with early stopping after 10 epochs of no improvement, using an effective batch size of 32 across four GPUs. The learning rate was set to 1× 10^−4^ with a weight decay of 1 ×10^−2^. Training was repeated three times with different random seeds for each split strategy, resulting in six runs per dataset and a total of 252 finetuned models. The best model for each run was selected using the epoch with the highest evaluation *R*^2^. All training was performed on NVIDIA A100 80-GB GPUs using PyTorch (89).

### Relative variability of site means and fraction of highly variable sites

The relative variability of site means (RVSM) is defined as the standard deviation of the site mean fitness values divided by the overall standard deviation. The site means are computed as

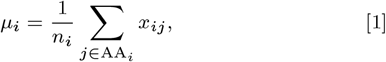

where *x*_*ij*_ is the fitness of amino acid *j* at site *i*, AA_*i*_ is the set of amino acids whose fitness has been measured at site *i*, and *n*_*i*_ is the number of elements in AA_*i*_. The RVSM is then given by

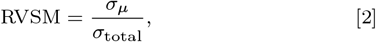

where *σ*_*µ*_ is the standard deviation of the site means and *σ*_total_ is the standard deviation of all fitness measurements in the dataset.

The fraction of highly variable sites (FHVS) is defined as the proportion of sites whose within-site standard deviation exceeds 70% of the total variability in the dataset. It is computed via the following equation:

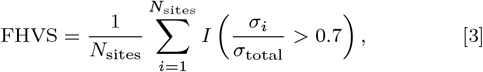

where *N*_sites_ is the total number of sites or residues in a protein, *σ*_*i*_ is the within-site standard deviation for site *i, σ*_total_ is the overall standard deviation of all fitness measurements in the dataset, and *I*(·) is the indicator function, equal to 1 when the condition inside the parentheses is true and 0 otherwise.

### Predicting model performance in ProteinGym data

We predicted model performance in ProteinGym data with a simple linear model with two predictors:

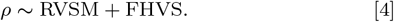

Here, *ρ* is the Spearman correlation coefficient reported by ProteinGym, and RVSM and FHVS are the relative variability of site means and the fraction of highly variable sites, respectively.

## Code and data availability

All the code used to generate embeddings, regression, finetuned models, data and results are available at: https://github.com/ziul-bio/DMS-SiteEffect-PLM

## ACKNOWLEDGMENTS

This work was supported by National Institutes of Health grants R01 AI169462 and R56 AI179799. The Texas Advanced Computing Center (TACC) and the NVIDIA Academic Grant program provided high-performance computing support. C.O.W. also acknowledges support from the Blumberg Centennial Professorship in Molecular Evolution.

## Supporting Information

**Fig. S1.**
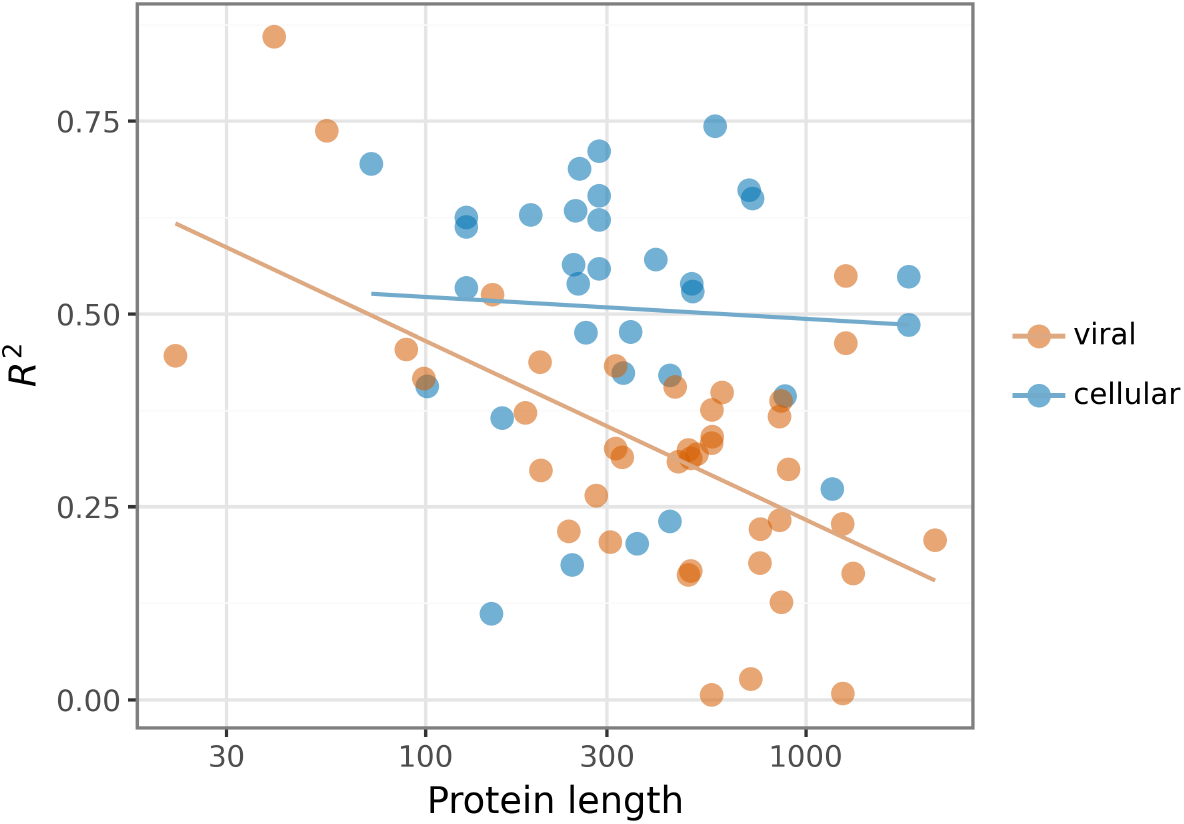
Effect of sample source and protein length on protein fitness landscape prediction. Transfer learning results using feature extraction from ESM C 600M. Each point corresponds to a dataset, colored blue for cellular and orange for viral. The y-axis reports the average *R*^2^ across five independent train–test splits, the x-axis shows protein length, and all results are from the pooled split.

**Fig. S2.**
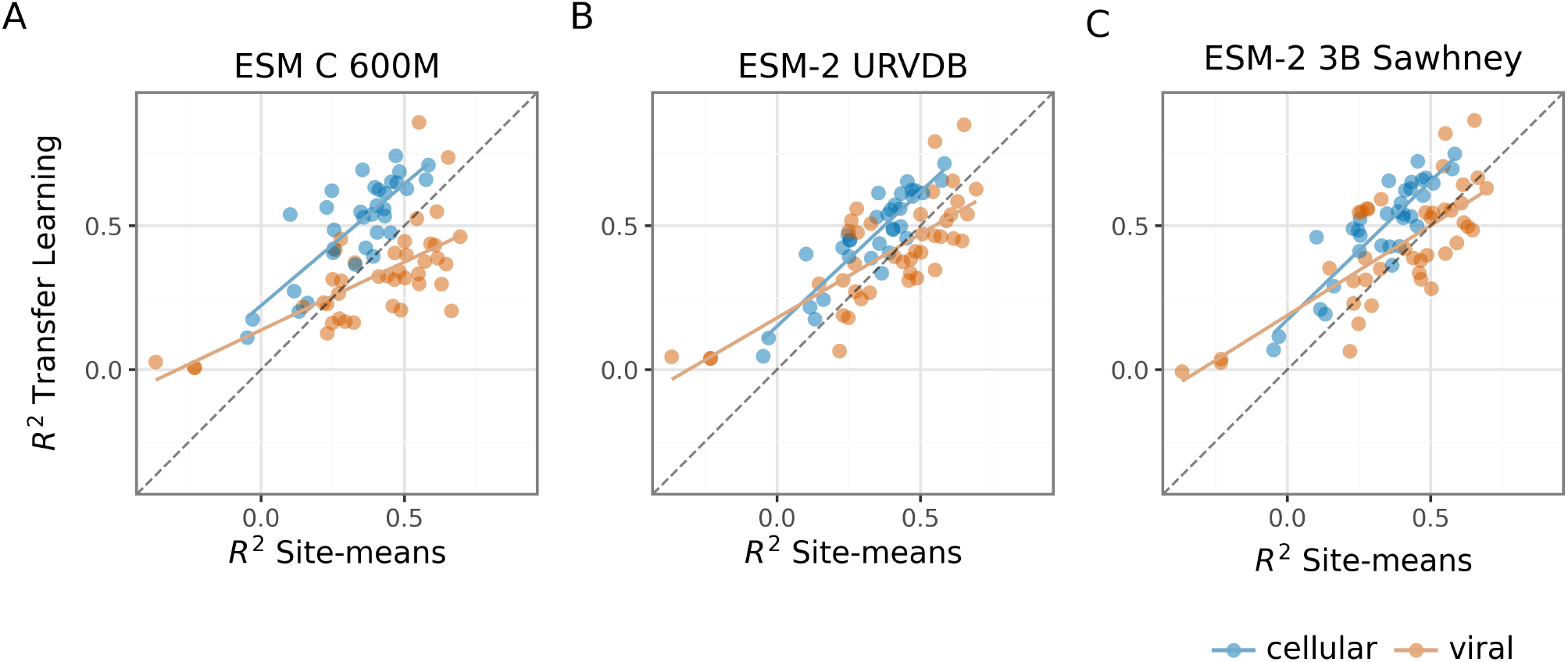
Site effects are observed across most datasets for both baseline and domain-adapted pLMs. The *R*^2^ achieved under transfer learning (Lasso regression, pooled data splits) is compared to the *R*^2^ obtained by simply taking the mean at each site in the training data as the prediction for the test data. The dashed line represents the one–one line where the transfer learning and site-means strategies perform equally. (A) Results for baseline model ESM C 600M. (B) Results for domain-adapted model ESM-2 650M URVDB (C) Results for domain-adapted model ESM-2 3B Sawhney.

**Fig. S3.**
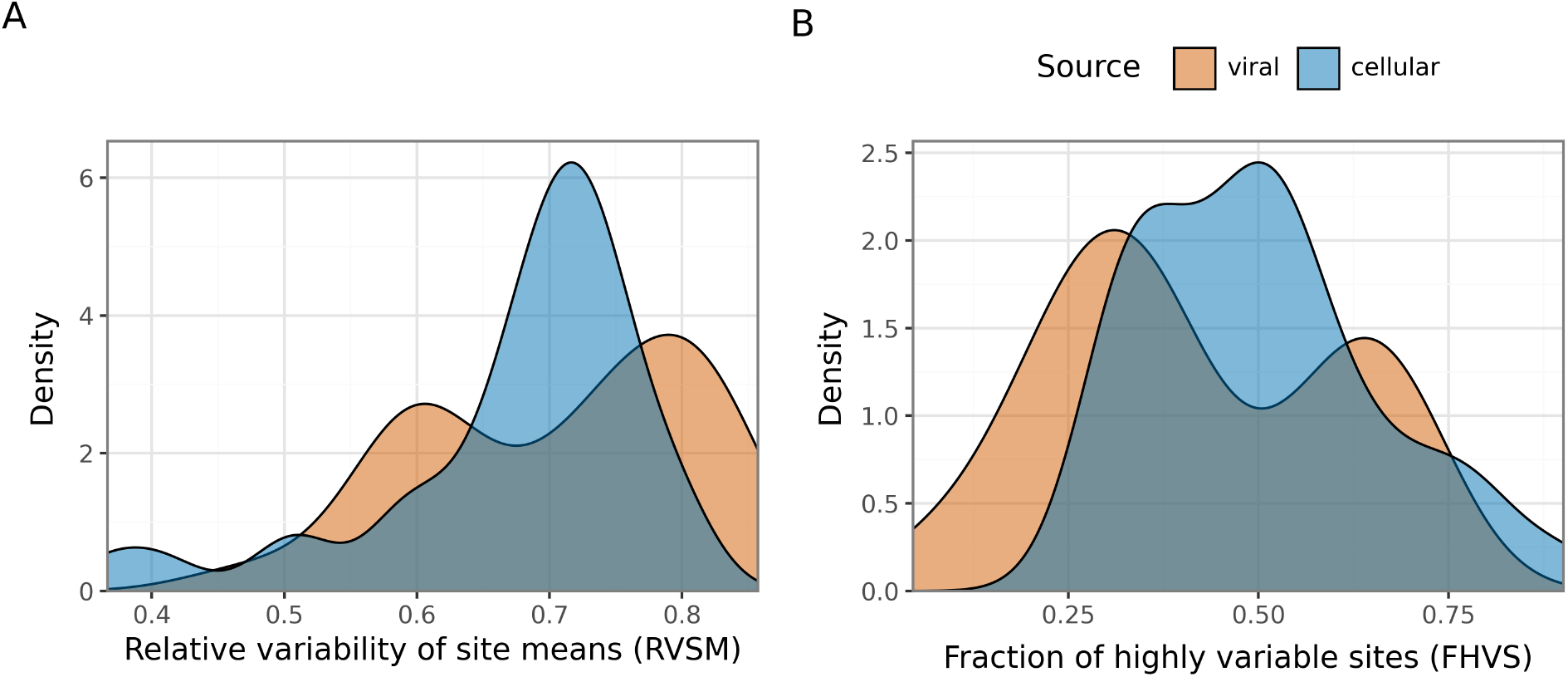
Site variability shift between viral and cellular datasets. (A) Density distribution of the relative variability of site means (RVSM) for each dataset. The x-axis shows the RVSM values within datasets. (B) Density distribution of the fraction of highly variable sites (FHVS) for each dataset, with the x-axis showing the FHVS values within datasets. In all cases the colors indicating data source (viral in orange, cellular in blue).

**Fig. S4.**
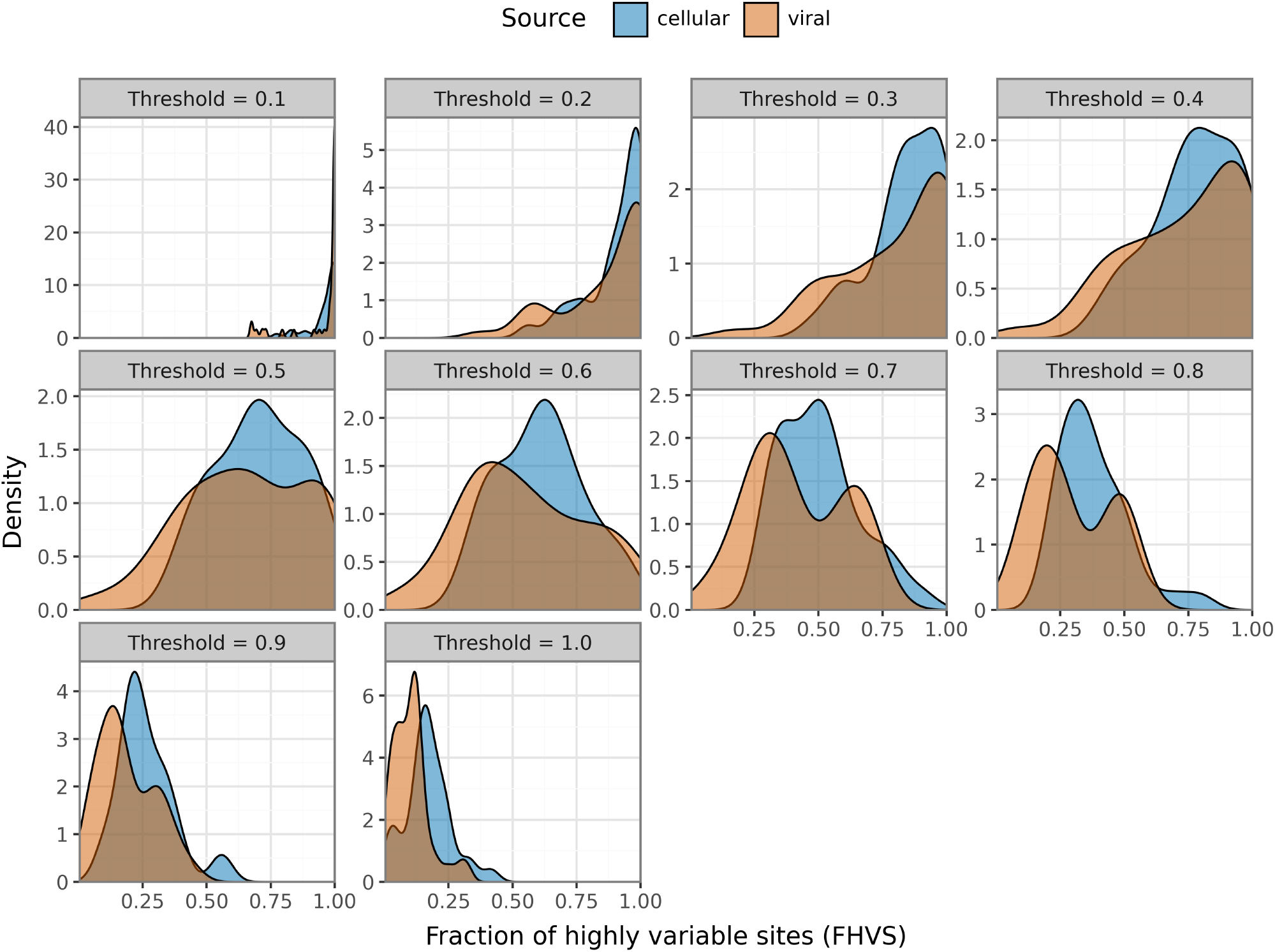
Distribution of the fraction of highly variable sites (FHVS) across thresholds. Density distributions of FHVS values are shown separately for viral and cellular datasets at different thresholds ranging from 0.1 to 1.0. Low thresholds produced highly compressed distributions in which most sites were classified as highly variable (thresholds from 0.1 to 0.4), whereas high thresholds yielded sparse distributions dominated by low FHVS values (thresholds 0.9 and 1.0). Intermediate thresholds, particularly near 0.7, provided the greatest discriminatory power, revealing the clearest separation between viral and cellular datasets and a pronounced bimodal distribution among viral proteins. At this threshold, approximately 50% of the cellular datasets exhibited high FHVS values, while viral datasets separated into distinct high-diversity and low-variability groups.

**Fig. S5.**
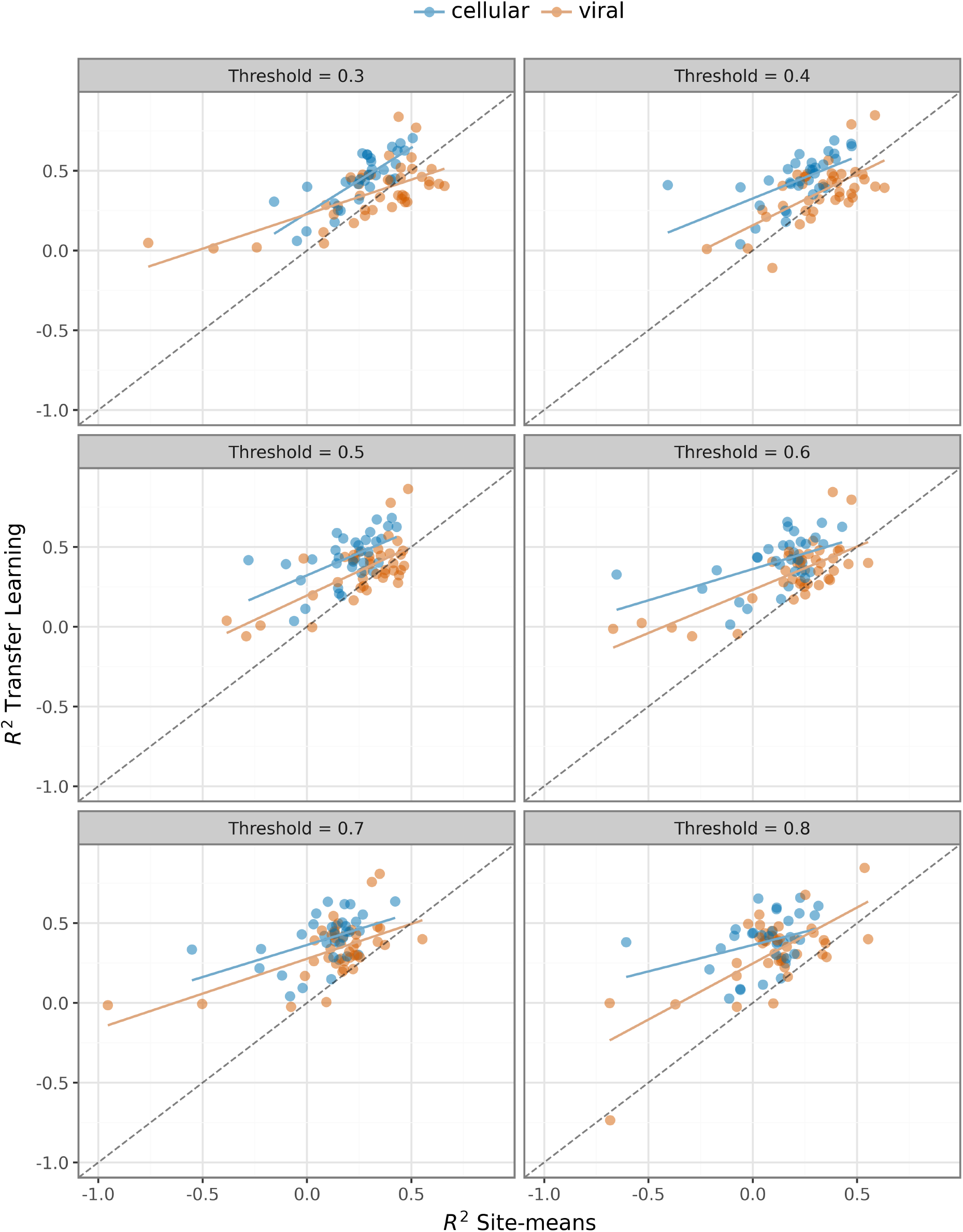
Effect of FHVS threshold choice on predictive performance correlations. The x-axis shows the predictive performance of site-means models, and the y-axis shows the performance of transfer learning using ESM-2 embeddings with Lasso regression. Each point represents a single DMS dataset, with cellular datasets shown in blue and viral datasets shown in orange. Across thresholds, the performance distributions of viral and cellular datasets progressively overlap, with the strongest convergence observed at thresholds near 0.7. The dashed diagonal line represents equal performance between the transfer learning and site-means approaches.

**Fig. S6.**
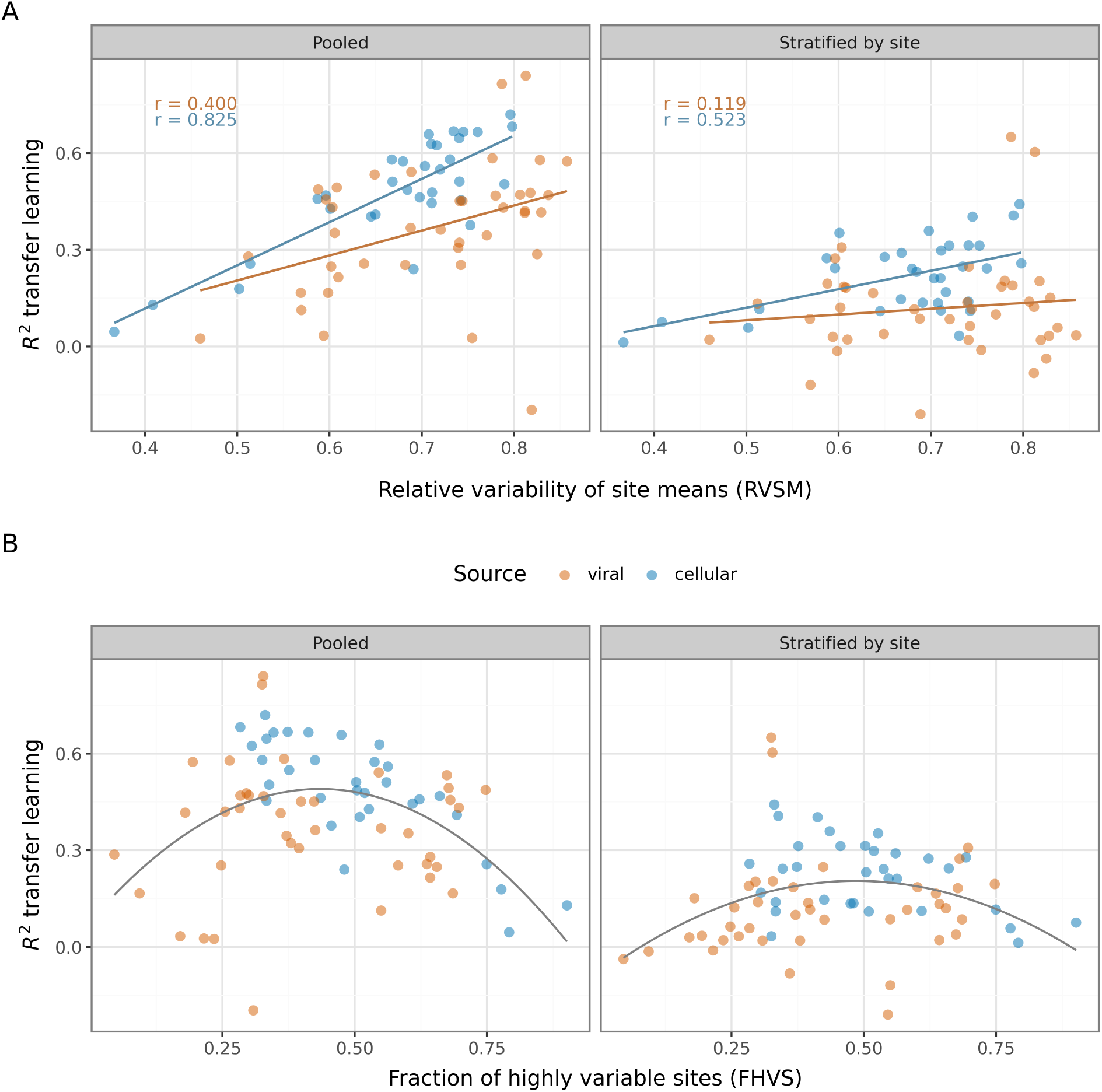
Impact of fitness variability measures on model performance, for different data splitting strategies. (A) Model performance (as measured by *R*^2^) as a function of the relative variability of site means (RVSM). The qualitative trend (*R*^2^ increases with larger RVSM) is preserved regardless of data splitting strategy. (B) Model performance (as measured by *R*^2^) as a function of the fraction of highly variable sites (FHVS). The qualitative trend (*R*^2^ is highest for intermediate values of FHVS) is preserved regardless of data splitting strategy. Results for both panels A and B were obtained with ESM-2 650M. Note that results for pooled splits are identical to those shown in Figure 2 and are reproduced here to facilitate comparison between the splitting strategies.

**Fig. S7.**
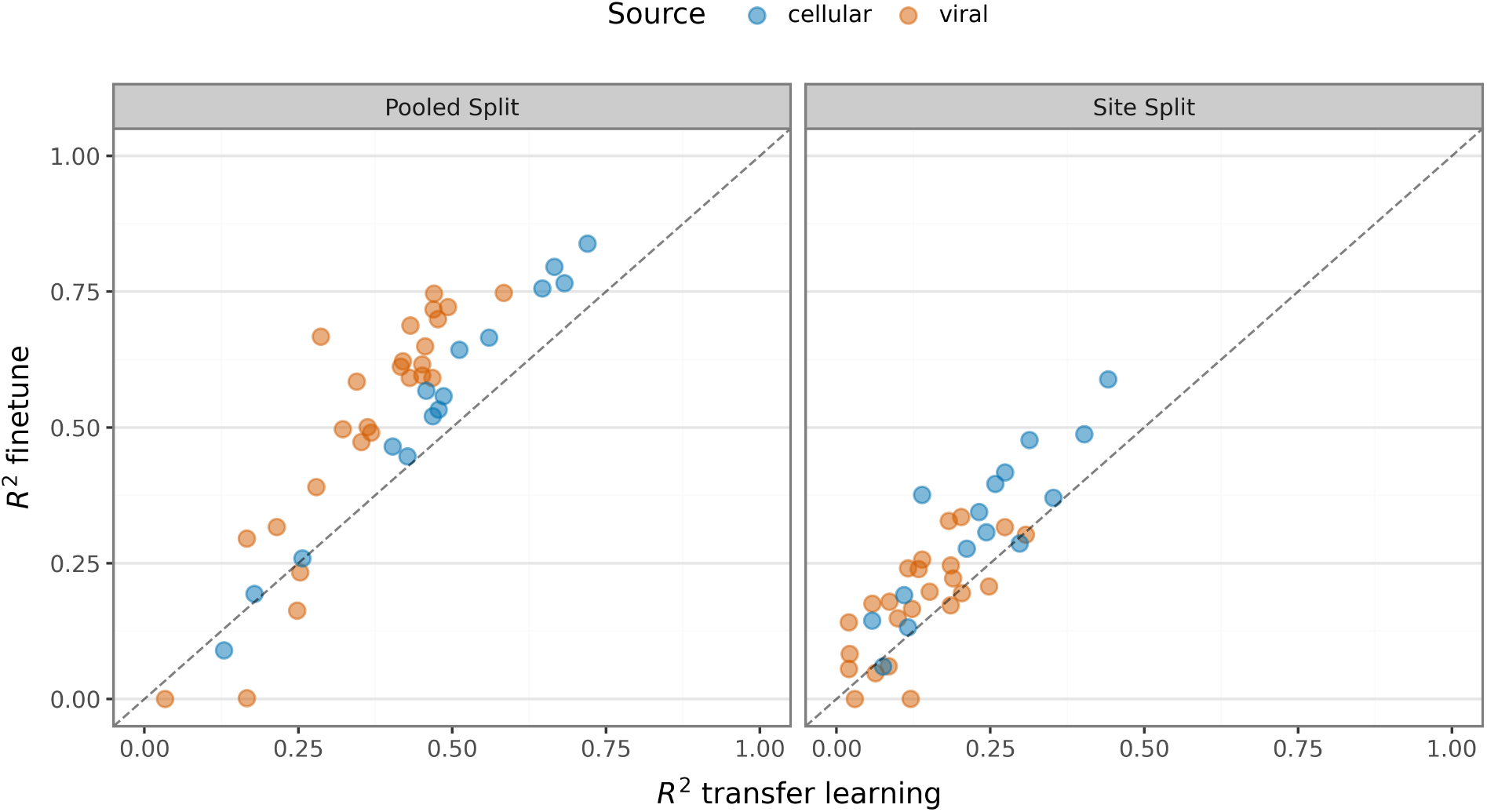
Comparing transfer learning strategies. Results from finetuning ESM-2 650M with LoRA are compared to transfer learning via feature extraction followed by a Lasso regression model. The Y-axis shows the average *R*^2^ from three independent train–test splits for finetuning, and the X-axis shows the average *R*^2^ from five independent train–test splits for the Lasso model. Facets indicate the split strategy, pooled on the left and site-stratified on the right. Colors show the data source, viral in orange and cellular in blue. Each point represents one of the 42 datasets, 15 cellular and 27 viral, see Methods for the selection criteria.).

**Fig. S8.**
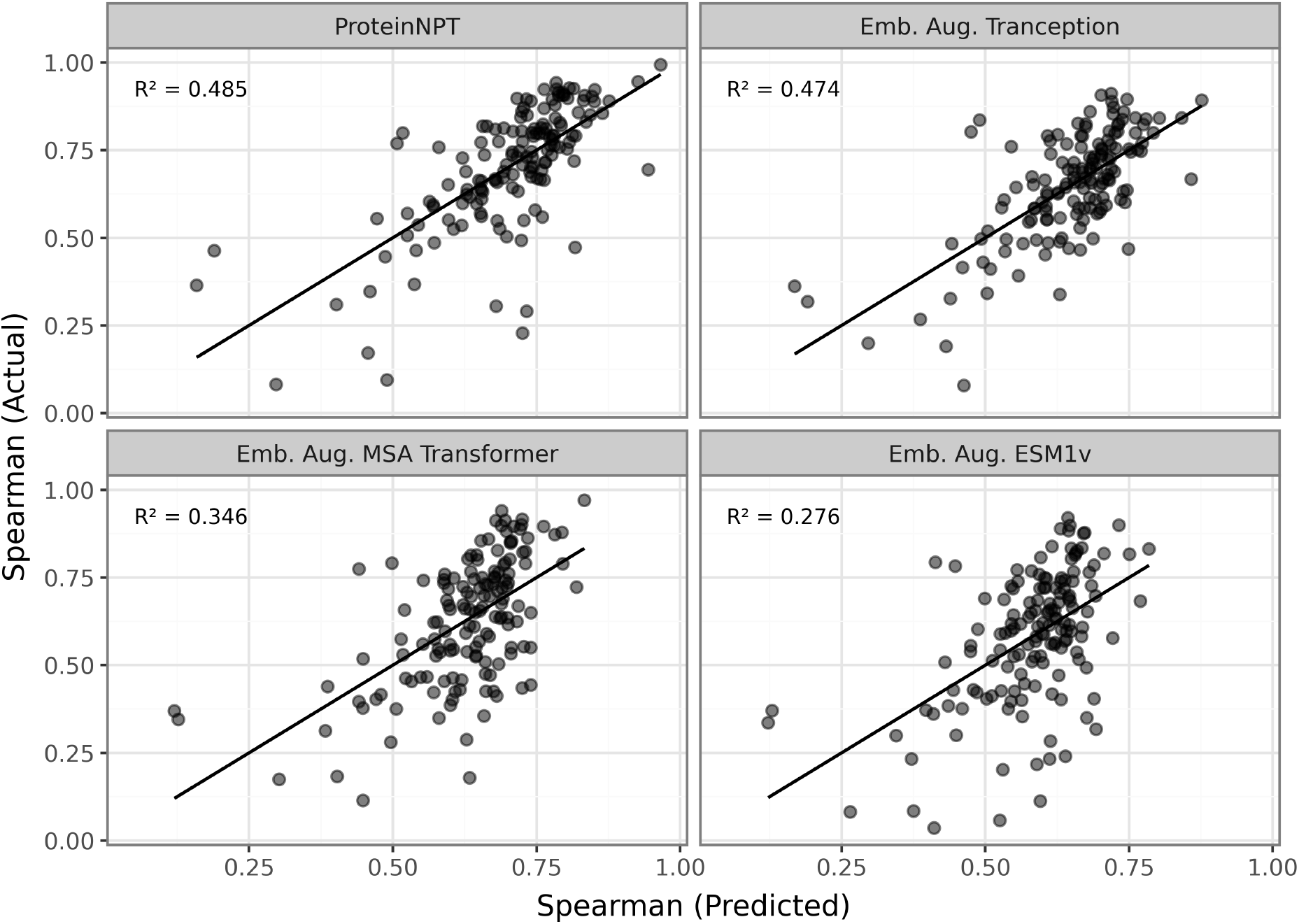
Predicting ProteinGyn dataset performance with variability metrics. Each point represents a Protein Gym dataset, showing actual versus predicted Spearman’s scores and each panel represent the second to fifth best models in ProteinGym. The top-left annotation reports the *R*^2^ from the ordinary least squares (OLS) model, summarizing how well the variability metrics explain performance across datasets. The reported results are obtain from the pooled split strategy.

**Fig. S9.**
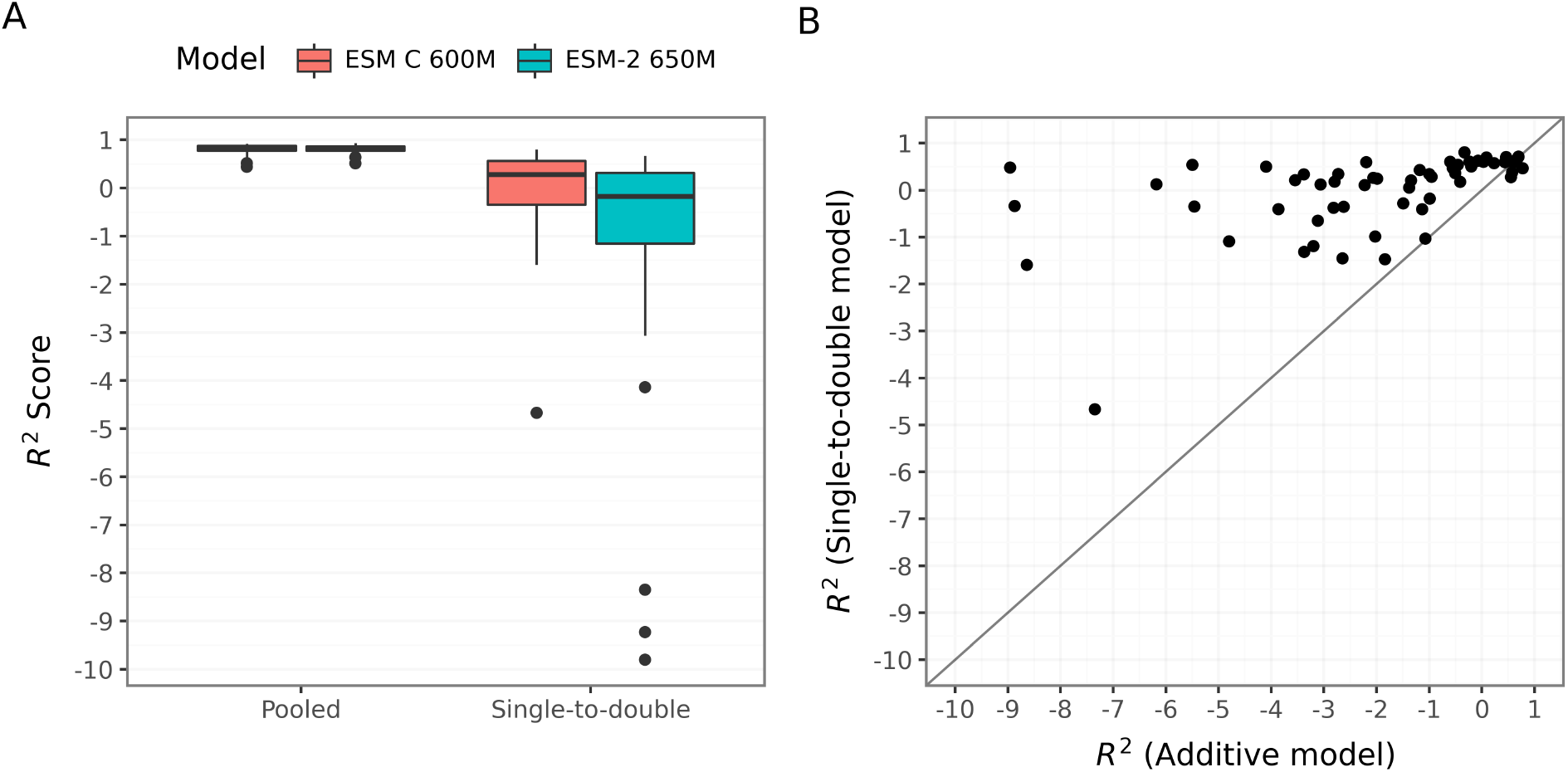
Evaluation of regression models trained on single-mutant DMS data for predicting double-mutant fitness effects across 58 datasets. (A) Distribution of *R*^2^ scores obtained under two evaluation strategies: pooled split, in which all variants were randomly divided into training and test sets, and single-to-double, in which models were trained exclusively on single mutants and evaluated on double mutants. The pooled split produced artificially high performance, consistent with information leakage arising from shared site-specific information between the training and test sets for both models (ESM C, red; ESM-2, cyan). By contrast, models trained on single mutants showed poor generalization to double mutants. (Large negative *R*^2^ values arise because the single-to-double models are highly biased and systematically mispredict pairwise fitness effects.) (B) Comparison between the additive baseline, in which double-mutant fitness was estimated as the sum of the corresponding single-mutant effects, and the single-to-double embedding-based model using ESM C 600M. Each dot represents one dataset. While the embedding-based model performs much better than the additive model, in absolute terms the embedding-based model still performs poorly. For most datasets the *R*^2^ is near or below zero, indicating limited predictive power and suggesting that representations learned from single mutants alone are insufficient to capture the higher-order epistatic interactions that shape the fitness of combinatorial variants.

## Notes

https://github.com/ziul-bio/DMS-SiteEffect-PLM

### Competing Interest Statement

The authors have declared no competing interest.

### Summary of Updates

Added a sensitivity analysis for the threshold choice in FHVS metric, added an analysis of double mutants, extended the Discussion.

## References

1. A Elnaggar, et al., Prottrans: Toward understanding the language of life through self-supervised learning. IEEE (TPAMI) 44, 7112–7127 (2021).

2. M Heinzinger, et al., Modeling aspects of the language of life through transfer-learning protein sequences. BMC Bioinform. 20, 723 (2019).

3. T Bepler, B Berger, Learning the protein language: Evolution, structure, and function. Cell Syst. 12, 654–669 (2019).

4. A Rives, et al., Biological structure and function emerge from scaling unsupervised learning to 250 million protein sequences. PNAS 118, e2016239118 (2021).

5. J Devlin, M Chang, K Lee, K Toutanova, Bert: Pre-training of deep bidirectional transformers for language understanding in NAACL HLT. Vol. 1, pp. 4171–4186 (2019).

6. A Villegas-Morcillo, et al., Unsupervised protein embeddings outperform hand-crafted sequence and structure features at predicting molecular function. Bioinformatics 37, 162–170 (2021).

7. EC Alley, G Khimulya, S Biswas, M AlQuraishi, GM Church, Unified rational protein engineering with sequence-based deep representation learning. Nat. Methods 16, 1315–1322 (2019).

8. Z Lin, et al., Evolutionary-scale prediction of atomic-level protein structure with a language model. Science 379, 1123–1130 (2023).

9. E Fenoy, AA Edera, G Stegmayer, Transfer learning in proteins: evaluating novel protein learned representations for bioinformatics tasks. Brief. Bioinform. 23, bbac232 (2022).

10. T Bepler, B Berger, Learning protein sequence embeddings using information from structure (arXiv preprint arXiv:1902.08661) (2019).

11. M Littmann, M Heinzinger, C Dallago, T Olenyi, B Rost, Embeddings from deep learning transfer GO annotations beyond homology. Sci. Reports 11, 1160 (2021).

12. T Hoffbauer, B Strodel, TransMEP: transfer learning on large protein language models to predict mutation effects of proteins from a small known dataset (bioRxiv) (2024) doi:10.1101/2024.01.12.575432.

13. H Dieckhaus, M Brocidiacono, NZ Randolph, B Kuhlman, Transfer learning to leverage larger datasets for improved prediction of protein stability changes. PNAS 121, e2314853121 (2024).

14. R Singh, et al., Learning the language of antibody hypervariability. PNAS 122, e2418918121 (2025).

15. JL Watson, et al., De novo design of protein structure and function with RFdiffusion. Nature 620, 1089–1100 (2023).

16. J Jumper, et al., Highly accurate protein structure prediction with AlphaFold. Nature 596, 583–589 (2021).

17. T Hayes, et al., Simulating 500 million years of evolution with a language model. Science 387, 850–858 (2025).

18. S Hayat, C Sander, DS Marks, A Elofsson, All-atom 3D structure prediction of transmembrane β-barrel proteins from sequences. PNAS 112, 5413–5418 (2015).

19. V Zambaldi, et al., De novo design of high-affinity protein binders with AlphaProteo (arXiv preprint arXiv:2409.08022) (2024).

20. S Bhat, et al., De novo design of peptide binders to conformationally diverse targets with contrastive language modeling. Sci. Adv. 11, eadr8638 (2025).

21. R Singh, S Sledzieski, B Bryson, L Cowen, B Berger, Contrastive learning in protein language space predicts interactions between drugs and protein targets. PNAS 120, e2220778120 (2023).

22. Z Wu, SBJ Kan, RD Lewis, BJ Wittmann, FH Arnold, Machine learning-assisted directed protein evolution with combinatorial libraries. PNAS 116, 8852–8858 (2019).

23. J Meier, et al., Language models enable zero-shot prediction of the effects of mutations on protein function. NeurIPS 34, 29287–29303 (2021).

24. N Brandes, G Goldman, CH Wang, CJ Ye, V Ntranos, Genome-wide prediction of disease variant effects with a deep protein language model. Nat. Genet. 55, 1512–1522 (2023).

25. JR Randall, LC Vieira, CO Wilke, BW Davies, Deep mutational scanning and machine learning for the analysis of antimicrobial-peptide features driving membrane selectivity. Nat. Biomed. Eng 8, 1–12 (2024).

26. M Huot, D Wang, J Liu, EI Shakhnovich, Predicting high-fitness viral protein variants with bayesian active learning and biophysics. PNAS 122, e2503742122 (2025).

27. DM Fowler, S Fields, Deep mutational scanning: a new style of protein science. Nat. Methods 11, 801–807 (2014).

28. A Nisthal, CY Wang, ML Ary, SL Mayo, Protein stability engineering insights revealed by domain-wide comprehensive mutagenesis. PNAS 116, 16367–16377 (2019).

29. EE Wrenbeck, LR Azouz, TA Whitehead, Single-mutation fitness landscapes for an enzyme on multiple substrates reveal specificity is globally encoded. Nat. Commun. 8, 15695 (2017).

30. RN McLaughlin, Jr., FJ Poelwijk, A Raman, WS Gosal, R Ranganathan, The spatial architecture of protein function and adaptation. Nature 491, 138–142 (2012).

31. BJ Livesey, JA Marsh, Updated benchmarking of variant effect predictors using deep mutational scanning. Mol. Syst. Biol. 19, e11474 (2023).

32. HY Kim, D Kim, Prediction of mutation effects using a deep temporal convolutional network. Bioinformatics 36, 2047–2052 (2020).

33. AJ Riesselman, JB Ingraham, DS Marks, Deep generative models of genetic variation capture the effects of mutations. Nat. Methods 15, 816–822 (2018).

34. S Gurev, N Youssef, N Jain, DS Marks, Variant effect prediction with reliability estimation across priority viruses (bioRxiv) (2025) doi:10.1101/2025.08.04.668549.

35. LC Vieira, ML Handojo, CO Wilke, Medium-sized protein language models perform well at transfer learning on realistic datasets. Sci. Reports 15, 21400 (2025).

36. S Gurev, N Youssef, N Jain, DS Marks, Sequence-based protein models for the prediction of mutations across priority viruses (ICLR 2025 Workshop GEM) (2025) https://openreview.net/forum?id=DvC6VL7TJK.

37. BJ Livesey, JA Marsh, Using deep mutational scanning to benchmark variant effect predictors and identify disease mutations. Mol. Syst. Biol. 16, e9380 (2020).

38. C Hsu, H Nisonoff, C Fannjiang, J Listgarten, Learning protein fitness models from evolutionary and assay-labeled data. Nat. Biotechnol. 40, 1114–1122 (2022).

39. CL Eng, Jc T., TW Tan, Predicting host tropism of influenza A virus proteins using random forest. BMC Med. Genomics 7, S1 (2014).

40. G Franzo, et al., Evaluation of different machine learning approaches to predict antigenic distance among Newcastle disease virus (NDV) strains. Viruses 17, 567 (2025).

41. MA Zeller, et al., Machine learning prediction and experimental validation of antigenic drift in H3 influenza A viruses in swine. Msphere 6, 10–1128 (2021).

42. SNH Bukhari, J Webber, A Mehbodniya, Decision tree based ensemble machine learning model for the prediction of Zika virus T-cell epitopes as potential vaccine candidates. Sci. Reports 12, 7810 (2022).

43. D Zhu, et al., Optimal trade-off control in machine learning–based library design, with application to adeno-associated virus (AAV) for gene therapy. Sci. Adv. 10, eadj3786 (2024).

44. G Passi, S Amittai, D Schneidman-Duhovny, Calibrated variant effect prediction at the residue level using conditional score distributions (bioRxiv) (2025) doi:10.1101/2025.11.24.690189.

45. ESM-Team, ESM Cambrian: Revealing the mysteries of proteins with unsupervised learning. (Online: https://evolutionaryscale.ai/blog/esm-cambrian) (2024).

46. T Bigot, S Temmam, P Pérot, M Eloit RVDB-prot, a reference viral protein database and its hmm profiles. F1000Research 8, 530 (2020).

47. R Sawhney, et al., Fine-tuning protein language models unlocks the potential of underrepresented viral proteomes. PeerJ 13, e19919 (2025).

48. L Trgovec-Greif, et al., VOGDB-database of virus orthologous groups. Viruses 16, 1191 (2024).

49. R Schmirler, M Heinzinger, B Rost, Fine-tuning protein language models boosts predictions across diverse tasks. Nat. Commun. 15 (2024).

50. P Notin, et al., ProteinGym: Large-scale benchmarks for protein fitness prediction and design. NeurIPS 36, 64331–64379 (2023).

51. P Notin, R Weitzman, D Marks, Y Gal, ProteinNPT: Improving protein property prediction and design with non-parametric transformers. NeurIPS 36, 33529–33563 (2023).

52. PM Groth, M Kerrn, L Olsen, J Salomon, W Boomsma, Kermut: Composite kernel regression for protein variant effects. NeurIPS 37, 29514–29565 (2024).

53. R Rao, et al., MSA Transformer (bioRxiv) (2021) 10.1101/2021.02.12.430858.

54. P Notin, et al., Tranception: protein fitness prediction with autoregressive transformers and inference-time retrieval in ICML. (PMLR), pp. 16990–17017 (2022).

55. C Hsu, H Nisonoff, C Fannjiang, J Listgarten, Learning protein fitness models from evolutionary and assay-labeled data. Nat. Biotechnol. 40, 1114–1122 (2022).

56. H Sun, et al., Accelerating protein engineering with fitness landscape modelling and reinforcement learning. Nat. Mach. Intell. 7, 1446–1460 (2025).

57. C Hsu, H Nisonoff, C Fannjiang, J Listgarten, Combining evolutionary and assay-labelled data for protein fitness prediction (bioRxiv) (2021) 10.1101/2021.03.28.437402.

58. Z Zhang, et al., A protein fitness predictive framework based on feature combination and intelligent searching. Protein Sci. 33, e5211 (2024).

59. M Li, et al., Sesnet: sequence-structure feature-integrated deep learning method for data-efficient protein engineering. J. Cheminformatics 15, 12 (2023).

60. S Gelman, SA Fahlberg, P Heinzelman, PA Romero, A Gitter, Neural networks to learn protein sequence–function relationships from deep mutational scanning data. PNAS 118, e2104878118 (2021).

61. Y Tan, R Wang, B Wu, L Hong, B Zhou, From high-throughput evaluation to wet-lab studies: advancing mutation effect prediction with a retrieval-enhanced model. Bioinformatics 41, i401–i409 (2025).

62. H Song, BJ Bremer, EC Hinds, G Raskutti, PA Romero, Inferring protein sequence-function relationships with large-scale positive-unlabeled learning. Cell systems 12, 92–101 (2021).

63. C Sruthi, M Prakash, Deep2Full: Evaluating strategies for selecting the minimal mutational experiments for optimal computational predictions of deep mutational scan outcomes. PLOS One 15, e0227621 (2020).

64. JA Barbero-Aparicio, A Olivares-Gil, J. Rodríguez, C García-Osorio, JF Díez-Pastor, Addressing data scarcity in protein fitness landscape analysis: A study on semi-supervised and deep transfer learning techniques. Inf. Fusion 102, 102035 (2024).

65. S Biswas, G Khimulya, EC Alley, KM Esvelt, GM Church, Low-N protein engineering with data-efficient deep learning. Nat. Methods 18, 389–396 (2021).

66. L Chen, et al., Learning protein fitness landscapes with deep mutational scanning data from multiple sources. Cell Syst. 14, 706–721 (2023).

67. Z Shamsi, M Chan, D Shukla, TLmutation: predicting the effects of mutations using transfer learning. The J. Phys. Chem. B 124, 3845–3854 (2020).

68. Y Xu, D Liu, H Gong, Improving the prediction of protein stability changes upon mutations by geometric learning and a pre-training strategy. Nat. Comput. Sci. 4, 840–850 (2024).

69. DJ Diaz, et al., Stability oracle: a structure-based graph-transformer framework for identifying stabilizing mutations. Nat. Commun. 15, 6170 (2024).

70. Z Zhang, et al., Protein language models learn evolutionary statistics of interacting sequence motifs. PNAS 121, e2406285121 (2024).

71. TN Starr, JW Thornton, Epistasis in protein evolution. Protein Sci. 25, 1204–1218 (2016).

72. M. González, LA Abriata, PE Tomatis, AJ Vila, Optimization of conformational dynamics in an epistatic evolutionary trajectory. Mol. Biol. Evol. 33, 1768–1776 (2016).

73. A Biswas, et al., Evolutionary sequence and structural basis for the epistatic origins of drug resistance in HIV (bioRxiv) (2025) doi:10.1101/2025.04.30.651576.

74. M Figliuzzi, H Jacquier, A Schug, O Tenaillon, M Weigt, Coevolutionary landscape inference and the context-dependence of mutations in β-lactamase TEM-1. Mol. Biol. Evol. 33, 268–280 (2016).

75. Y Sun, Y Shen, Structure-informed protein language models are robust predictors for variant effects. Hum. Genet. 144, 209–225 (2025).

76. R Sanjuán, P Domingo-Calap, Genetic diversity and evolution of viral populations. Encycl. virology 1, 53 (2021).

77. Y Li, S Arcos, KR Sabsay, AJW Te Velthuis, AS Lauring, Deep mutational scanning reveals the functional constraints and evolutionary potential of the influenza A virus PB1 protein. J. virology 97, e01329–23 (2023).

78. A Lauring, R Frydman, J. amd Andino, The role of mutational robustness in RNA virus evolution. Nat. Rev. Microbiol. 11, 327–336 (2013).

79. CL Worth, S Gong, TL Blundell, Structural and functional constraints in the evolution of protein families. Nat. Rev. Mol. Cell Biol. 10, 709–720 (2009).

80. PR Woolley, A Feller, AO Ellington, CO Wilke, Overestimating zero-shot fitness prediction: Broad benchmarks mask local failures and practical limitations (bioRxiv) (2026) 10.64898/2026.06.04.73012.

81. B Ding, H Qian, J Zhou, Activation functions and their characteristics in deep neural networks in CCDC. (IEEE), pp. 1836–1841 (2018).

82. A Babjac, et al., Adapting protein language models for explainable fine-grained evolutionary pattern discovery in BIBM. (IEEE), pp. 2609–2616 (2023).

83. Z Yang, et al., Self-distillation bridges distribution gap in language model fine-tuning (arXiv preprint arXiv:2402.13669v2) (2024).

84. AJ Riesselman, JB Ingraham, DS Marks, Deep generative models of genetic variation capture the effects of mutations. Nat. Methods 15, 816–822 (2018).

85. M McCloskey, NJ Cohen, Catastrophic interference in connectionist networks: The sequential learning problem. Psychol. Learn. Motiv. 24, 109–165 (1989).

86. R Tibshirani, Regression shrinkage and selection via the lasso. J. Royal Stat. Soc. Ser. B (Methodological) 58, 267–288 (1996).

87. S Mangrulkar, et al., PEFT: State-of-the-art parameter-efficient fine-tuning methods (Online: https://github.com/huggingface/peft) (2022).

88. EJ Hu, et al., Lora: Low-rank adaptation of large language models. ICLR 1, 3 (2022).

89. J Ansel, et al., Pytorch 2: Faster machine learning through dynamic python bytecode transformation and graph compilation in ACM. (2024).

